# TOMM40-APOE chimera linking Alzheimer’s highest risk genes: a new pathway for mitochondria regulation and APOE4 pathogenesis

**DOI:** 10.1101/2024.10.09.617477

**Authors:** Jinglei Xu, Jingqi Duan, Zhiqiang Cai, Chie Arai, Chao Di, Christopher C. Venters, Jian Xu, Maura Jones, Byung-Ran So, Gideon Dreyfuss

**Affiliations:** Department of Biochemistry and Biophysics University of Pennsylvania, Perelman School of Medicine Philadelphia, Pennsylvania 19104, USA

**Author notes:** Equal contributions (Jinglei Xu, Jingqi Duan).

## Abstract

The patho-mechanism of apolipoprotein variant, APOE4, the strongest genetic risk for late-onset Alzheimer’s disease (AD) and longevity, remains unclear. APOE’s neighboring gene, TOMM40 (mitochondria protein transport channel), is associated with brain trauma outcome and aging-related cognitive decline, however its role in AD APOE4-independently is controversial. We report that TOMM40 is prone to transcription readthrough into APOE that can generate spliced TOMM40-APOE mRNA chimera (termed T9A2) detected in human neurons and other cells and tissues. T9A2 translation tethers APOE (normal APOE3 or APOE4) to near-full-length TOM40 that is targeted to mitochondria. Importantly, T9A2-APOE3 boosts mitochondrial bioenergetic capacity and decreases oxidative stress significantly more than T9A2-APOE4 and APOE3, and lacking in APOE4. We describe detailed interactomes of these actors that may inform about the activities and roles in pathogenesis. T9A2 uncovers a new candidate pathway for mitochondria regulation and oxidative stress-protection that are impaired in APOE4 genotypes and could initiate neurodegeneration.

## Introduction

Despite intensive research, the pathogenic mechanism of late-onset Alzheimer’s disease (AD), the most common neurodegenerative disease and cause of dementia, remains poorly understood. Drug development efforts focused mainly on hallmarks of advanced AD, particularly amyloid-β (Aβ) plaques, have been largely unsuccessful, indicating that additional therapy targets are needed. APOE4, a variant of the apolipoprotein (APOE) is the highest genetic risk for AD and a major risk factor for other neurodegenerative diseases, atherosclerosis (*1–7*), and human longevity (*8*, *9*). APOE4 is not rare; approximately 25% of world population and the majority of AD patients carry at least one APOE4 allele, which increases AD risk 3-4-fold compared with the common non-pathogenic APOE3 and two APOE4 alleles increase AD risk 12-14-fold and lowers age of onset by up to 12 years (*7*). APOE is a secreted lipid-binding protein that forms cholesterol- and triglycerides-rich lipoprotein particles. It is produced mainly in the liver and in brain, primarily by astrocytes that supply and recycle lipids to neurons via lipoprotein receptors. APOE4 differs from APOE3 in a single amino acid, C112R, which alters APOE4 conformation and promotes multiple aspects of AD pathology, including accumulation of amyloid and neurofibrillary tangles (*10*), mitochondrial dysfunction (*11*), alters lipid metabolism (*12*, *13*), cell signaling, induces neuro-inflammation and causes neuronal death. Nevertheless, the mechanisms by which a bewildering myriad of triggers turn APOE4 from a risk factor to an actual disruptor of many processes in different cell types and across disease stages remain unknown (*14*). Some of the above-mentioned effects can be explained by extracellular APOE4. However, a number of studies point to a crucial role for APOE4 produced in neurons themselves in cell-autonomous death.

Neurons do not normally express APOE but turn it on when stressed or injured (*15–19*). Under oxidative stress, neurons produce and secrete APOE, which facilitates disposal of peroxidized lipids, generated by mitochondria reactive oxygen species (ROS) resulting from intensive neuronal activity or mitochondrial impairment, because neurons are not equipped to detoxify them (*20*). APOE4 is defective in such function (loss of function, LOF), which could lead to accumulation of damaging reactive intermediates (*21*). Another mechanism implicates stress-induced APOE4 as a direct cell autonomous agent of mitochondrial damage. Accordingly, a fraction of APOE escapes the secretory pathway from the endoplasmic reticulum (ER) or Golgi to the cytoplasm, where it is cleaved by as yet unidentified neuron-specific protease(s), generating fragments that bind to and are toxic for mitochondria (gain of function, GOF) (*17–19*). APOE3 does not produce such toxic fragments. This model is attractive because it places APOE4 upstream of mitochondrial dysfunction which significantly precedes appearance of amyloid plaques (*22*), thus implicating APOE4 (fragments) in initiating AD and other diseases. Yet, essential information about the escape route, proteolysis, mitochondrial targeting and mode of action of APOE4 fragments are lacking.

Mitochondria are essential for oxidative phosphorylation and metabolism, proteostasis, immune response, and apoptosis, and their dysfunction has been linked to all neurodegenerative diseases, and their numbers decrease in ageing (*23–28*). Interestingly, polymorphisms in APOE neighboring gene, TOMM40, encoding the translocase of the outer mitochondrial membrane (OMM), have also been identified from genome-wide association studies (GWAS) as high-risk gene for AD, poor outcome from head trauma and age-related cognitive decline (*29–32*). GWAS have also revealed strong linkage disequilibrium between APOE and TOMM40, however, in other studies TOMM40 SNPs could not be separated from APOE4, casting doubt about APOE4-independent role for TOMM40 in AD (*19*, *27*, *28*), which left TOMM40 largely ignored in relation to AD and the potential interaction between the two genes remains unexplained.

## Results

### TOMM40 transcription readthrough into APOE generates chimeric TOMM40-APOE splicing

From visual survey of RNA-seq reads from various experiments in our laboratory, we noticed frequent and robust increases of nascent RNA extending continuously downstream of TOMM40 through the intergenic region [2.1 kilobases (kb)] and into APOE, which is on the same DNA strand and sense orientation. For example, in HeLa and other human cells inhibition of U1 snRNP (U1), which has roles in splicing and telescripting (transcription elongation and termination control), with antisense morpholino oligonucleotide (U1 AMO) (*35*, *36*), or inhibition of RNA polymerase II phosphorylation by CDK12 with THZ531 (*37*) (**Fig. 1A and B**). Reads in APOE (all exons) increased several-fold (**Fig. 1D**). Reads over introns were also increased consistent with the effects of the inhibitors on splicing, although splicing was still evident in both genes under these experimental conditions. The corresponding controls, non-targeting AMO (cAMO) or DMSO had no effect. RT-PCR using primers in TOMM40 last exon (exon10; T10) and APOE first exon (exon1; A1) in U1 AMO (**Fig. 1C**) showed uninterrupted transcripts, representing transcription readthrough (hereafter, readthrough) from TOMM40 into APOE, and suggesting that at least some of the transcription in APOE originated from TOMM40 instead of or in addition to *de novo* transcription from APOE transcription start site (TSS). Like most cells, including neurons, HeLa do not normally express APOE or only very little.

**Fig. 1.**
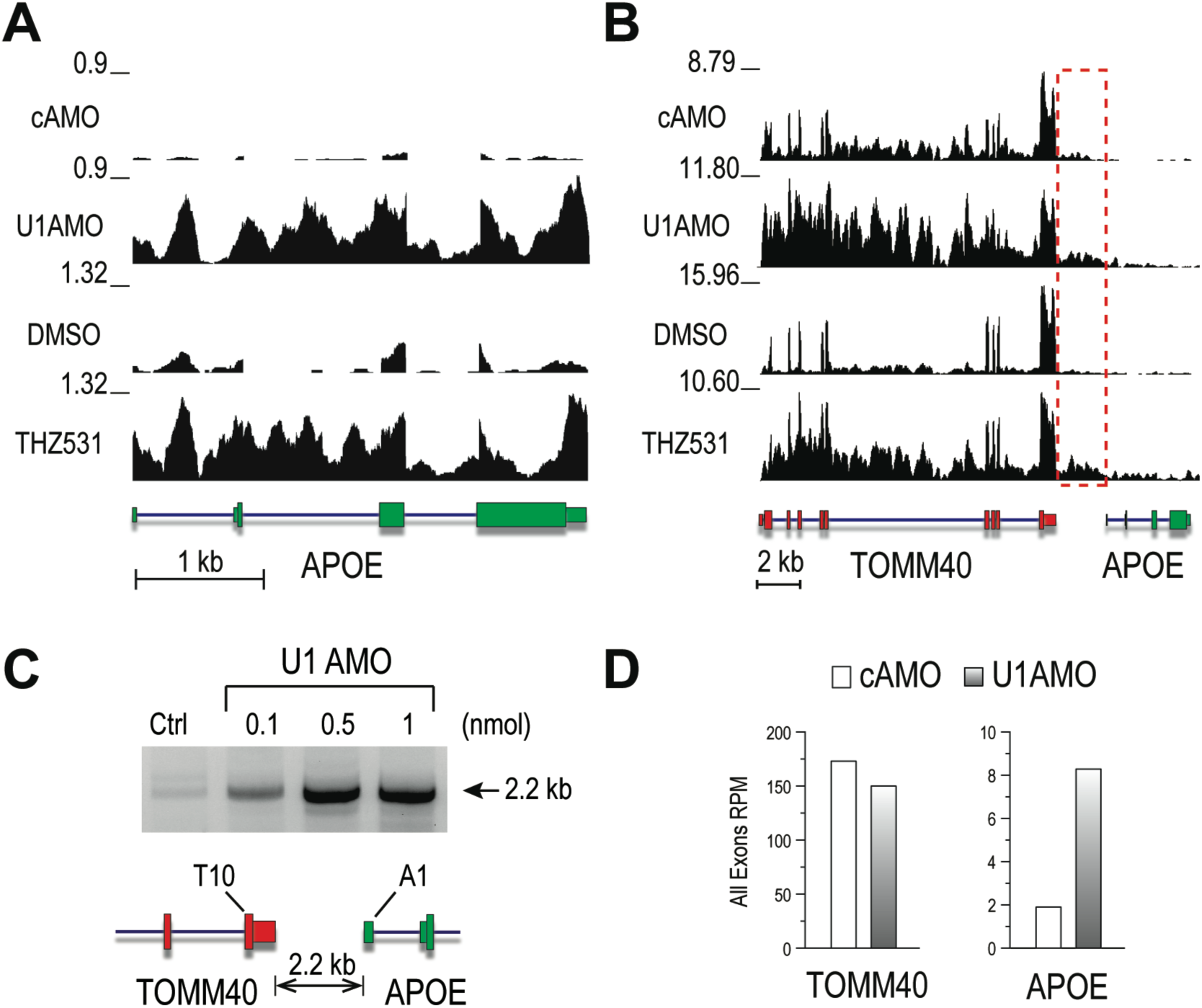
Transcription readthrough from TOMM40 to APOE upon inhibition of U1 or CDK12. (**A**) RNA-seq reads of nascent transcripts, labeled for 30min with 4-thiouridine from HeLa cells at 6h post-transfection with cAMO or U1 AMO, and DMSO control and the CDK12 inhibitor THZ531 mapped to APOE. The Y axis shows the maximal value of reads normalized for sequencing depth (reads per million, RPM). For clarity, the control RPMs are plotted on the scale of U1 AMO and THZ531, respectively. The gene structure according to NCBI RefSeq is shown below the X-axis. Exons are represented by boxes, mRNA untranslated regions (UTRs) as thinner boxes, and introns as connecting lines. (**B**) Mapped reads from the same cells, RNA labeling and RNA-seq protocol as in **A** for the TOMM40-APOE genomic region (hg38, chr19:44,891,220-44,909,393). The Y-axis shows the maximal RPM values of each. The gene structures are shown below the X-axis. The dashed red box indicates the intergenic region. (**C**). RT-PCR using primers in TOMM40 exons10 (T10) and APOE exon1 (A1), respectively, as indicated. The RT-PCR primers are 2.2 kb apart at the indicated positions on the gene structure. (**D**). Quantitation of RPMs in all exons of TOMM40 and APOE, showing a strong (>4-fold) increase in APOE expression due to readthrough from TOMM40.

Widespread readthrough occurs from >1,000 genes under stress conditions, including hyperosmotic shock, hypoxia, oxidative stress and infections with HSV-1 and influenza virus (*38*– *41*). While the molecular mechanisms and biological functions of readthrough are unclear, readthrough-prone genes generally have non-canonical and weaker transcription-terminating polyadenylation signal (PAS) hexanucleotide, instead of a canonical AAUAAA (*38*, *42*). TOMM40 also has a non-canonical PAS (AAUAUA), unlike APOE.

### TOMM40-APOE spliced chimera (T9A2) expressed in human cells and tissues

Notably, in many of our samples, we observed splicing in the TOMM40-APOE readthrough transcripts albeit at low levels, primarily from TOMM40 exon9 (the 5’ss of intron9) to APOE exon2 (the 3’ss of intron1), namely, alternatively spliced TOM40Δexon10-APOEΔexon1, which we termed T9A2. Less abundant isoforms, e.g., TOM40Δexon10-APOEΔexon1,2, (T9A3) were also detected. The potential importance of TOMM40-APOE chimera motivated us to ask if the signatory T9A2 RNA is expressed in any human cells and tissues. Not knowing if or where these may be found, we devised a protocol to search thousands of publicly accessible RNA-seq runs (called SRRs) at the NCBI Sequence Read Archive (SRA) in parallel with a 40nt sequence of the T9A2 splice junction, including 20nt upstream and downstream of the junction. Sequence similarity searches were performed using the BLAST algorithm (SRA Toolkit) at high stringency, requiring ≥90% identity, ≥75% query coverage, and Evalue ≤1E-6. BLAST of the human genome with the same sequence and settings gave no hits as expected for an RNA sequence generated post-transcriptionally by splicing. SRRs were selected for searches based on keywords, e.g., brain, neuron, and various diseases and physiological conditions, and batch size for each BLAST search was set according to available computing resources. A search of nearly 5,000 SRRs (during 2020-2023), which represents a small fraction of the SRA, detected T9A2 in hundreds of SRRs from diverse cell types and tissues. Overall, 478 (∼10%) out of 4,761 SRRs from 114 studies (SRA BioProjects), had T9A2 hits, of which 78 had ≥3 reads. A complete list of the metadata from these BioProjects and including quantitation of T9A2 (numbers of hits and RPM) and of T6T7 and A2A3 other splice junctions in TOMM40 and APOE as proxies of expression of these genes and probe sequences are shown in supplementary **Table S1** and **Data S1-8.**

To verify the BLAST results we downloaded the raw RNA-seq from several studies that had ≥3 T9A2 hits in multiple samples and analyzed them using the same pipeline and settings as all the other RNA-seq in this manuscript. Data for representative studies, including scatter plots and sashimi plots, shown in **Fig. 2**, validate the BLAST searching scheme and show chimeric TOMM40-APOE mRNAs in many independent experiments and conditions. Several points of interest raised by these surveys include the possibility that T9A2 expression is increased under pathological and adverse conditions and regulated by genetic and environmental factors (**Fig. 2A**). Much larger studies will be needed to assess the significance of these observations.

**Fig. 2.**
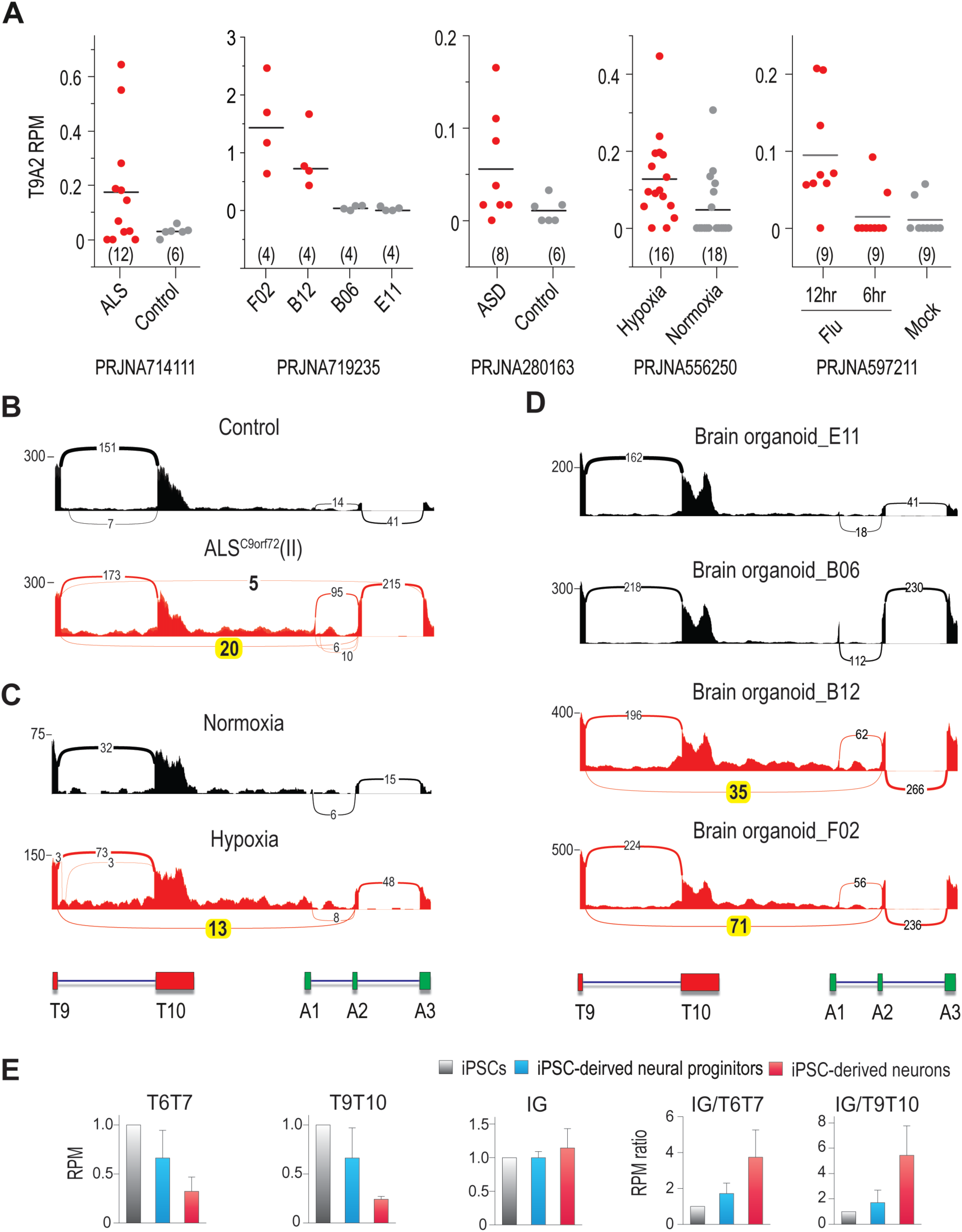
TOMM40 readthrough and TOMM40-APOE splicing (T9A2) in selected RNA-seq of human samples. (**A**). Scatter plots of RPM of BLAST detected T9A2 in SRRs of the indicated BioProjects. The datapoints from patient/experimental and the corresponding controls are shown in red and in gray, respectively. (**B-D**). Sashimi plots showing mapped RNA-seq reads and splice junctions in the region from TOMM40 exon9 (T9) to the end of APOE exon3 (A3). **B**. RNA-seq of iPSC derived motor neurons from control (upper, SRR13955538, SRR13955539, SRR13955540) and an ALS patient with C9orf72 mutation (lower, SRR13955547, SRR13955548, SRR13955549) in PRJNA714111. Participant i.d. was indicated in parenthesis. **C.** RNA-seq of neurons derived from human pluripotent stem cell treated with normoxia (upper, SRR9736783) and hypoxia (lower, SRR9736789) in PRJNA556250. **D.** RNA-seq of brain organoids derived from pathogenic tau-V337M mutation isogenic CRISPR-corrected lines (GIH7-F02, GIH6-E11, NDB06, and GIH7-B12) in PRJNA719235. (**E**). Histogram showing TOMM40 expression down-regulation with neuronal differentiation in PRJNA280163. Normalized RPM is the ratio of the indicated TOM40 mRNA splice junctions or intergenic reads (IG) in iPSC-derived neural progenitors (blue bars) or iPSC-derived neurons (red bars) to the same values in the iPSCs from which they were generated (gray). Splice junctions used as expression level proxies: T6T7: TOMM40 exon6-exon7 junctions; T9T10: TOMM40 exon9-exon10 junctions; IG: TOMM40-APOE intergenic region.

For examples, in iPSC-derived motor neurons (MNs) from amyotrophic lateral sclerosis (ALS) patients (BioProject PRJNA714111) (*43*), one of two patients (patient II) with pathogenic repeats expansion in C9orf72, had 3-fold higher T9A2 expression (RPM mean=0.49) compared to the cohort (RPM mean=0.13), including ALS patients with FUS mutations and healthy controls. Patient II also had higher TOMM40 intergenic/readthrough reads than the others (**Fig. 2B, Data S2).** In brain organoids from four individuals in a study of the role of frontotemporal dementia (FTD) microtubule-associated protein tau (*MAPT*) mutation (FTD-tau) (PRJNA719235) (*44*), high T9A2 reads were detected in multiple brain organoids differentiated from three FTD-tau carriers isogenic corrected (V337V) controls iPSC lines at a 2-month developmental stage (**Fig. 2D, Data S7**).

In iPSC-derived neurons from a study on cellular models for autism spectrum disorder (ASD) (PRJNA280163) (*45*) most of the T9A2 reads (10 out of 14 ASD and control) were from the autistic cohort (**Data S4**). T9A3 was also detected. In this study, which included iPSCs, iPSC-derived neuronal progenitors, and iPSC-derived neurons from each sample, most (9/13) T9A2 reads were from the iPSC-derived neurons compared with only 3/13 in iPSC and none from iPSC-derived neuronal progenitors. Interestingly, in the entire cohort, TOMM40 expression, reflected by T6T7 RPM, was significantly lower in iPSC-derived neurons compared with the other cell types (**Fig. 2E**). The reads ratio in readthrough vs T9T10 or vs T6T7 was also higher in iPSC-derived neurons, suggesting that a higher fraction of TOMM40 transcription becomes TOMM40-APOE, which may contribute to the lower TOMM40 expression in the iPSC-derived neurons, potentially making neurons more vulnerable to TOMM40 loss from readthrough. Readthrough was evident in many iPSC-derived neurons in all the studies examined, and T9A2 was also detected in multiple samples of iPSC- or hPSC-derived MNs (PRJNA556250) (*46*)(**Data S5**) in both soma and neurites, supporting the conclusion that T9A2 mRNAs are exported from the nuclei and transported to neurites.

T9A2 expression was generally higher under various cell stress conditions. For example, in hESC-derived soma, T9A2 average RPM was 0.21 in hypoxia versus 0 in normoxia (**Fig. 2C** and **Data S5**). In Influenza A virus (IAV) infection of A549 lung epithelial cells (PRJNA597211) (**Data S6**) at 12h post-infection, there was robust TOMM40 readthrough. The T9A3 spliced isoform and splicing from a cryptic 5’ss in the 3’UTR was detected in IAV, consistent with splicing dysregulation in IAV infection (*47*) and in HSV-1 infection (PRJNA688309) (**Data S8**) (*48*) and supporting the conclusion that readthrough makes a TOMM40-APOE pre-mRNAs that increases under cell stress. T9A2 was also detected in post-mortem brains, such as frontal cortex and/or cerebellum from a 27-individuals cohort, including ALS patients with pathogenic C9orf72 mutations (cALS) or sporadic ALS (sALS), and non-ALS controls (healthy or other neuropsychiatric conditions) (PRJNA279249) (*49*). Out of 18 ALS patients, 10 (56%) had T9A2 or T9A3 compared with 3 out of 9 (33%) in non-ALS controls (**Data S3**). T9A2 or T9A3 expression levels in these brain samples were generally low (RPM ≤0.09; <10% of T9T10), however they were similar to T1T2 in the same samples. T9A2 level in expressing cells in brain could be much higher if it is limited to specific cell types, disease stages or transiently expressed.

### T9A2 has a translation open reading frame (ORF) for TOM40-APOE fusion protein

T9A2 mRNA has an ORF for TOM40 lacking the C-terminal 46aa encoded in T10 (TOM40ΔC46). T9 splices into the APOE 5’UTR, 25nt upstream of the APOE translation start. Remarkably, the 24nt of APOE exon2 5’UTR has the potential to encode an octapeptide (DWPITGRK; oct) in-frame with TOM40ΔC46 and the entire APOE coding sequence (**Fig. S1**). The predicted ORF suggests a fusion protein comprising APOE, including its N-terminal 18 amino acid secretory signal peptide (sec) linked to TOM40ΔC46 by the octapeptide. This novel T9A2 protein (TOM40ΔC46-oct-sec-APOE) is predicted to have 640aa and molecular mass of 70kDa. Unlike APOE, which has the sec removed as it translates into the endoplasmic reticulum (ER), in T9A2, it would be in the middle of the protein, and thus will likely translate in the cytoplasm and remain intracellular. T9A3 also has an ORF similar to T9A2, however it lacks the oct and first 14aa of the secretory sequence (**Fig. S1**).

To test if the predicted T9A2 ORF is translatable, we constructed cDNAs of APOE3, APOE4, TOMM40 and T9A2s with APOE3 and APOE4 variants with C-terminal Flag tag in tetracycline-inducible (tet-ON) lentivirus vector. Western blots of transduced HeLa showed a doxycycline (DOX) (a tetracycline analog) inducible ∼70kDa band, the expected T9A2 mass, reactive with antibodies to Flag, APOE and TOM40 (**Fig. 3A**). The expression levels of T9A2 were consistently much lower compared with the APOEs and TOMM40 constructs (**Fig. 3B**).

**Fig. 3.**
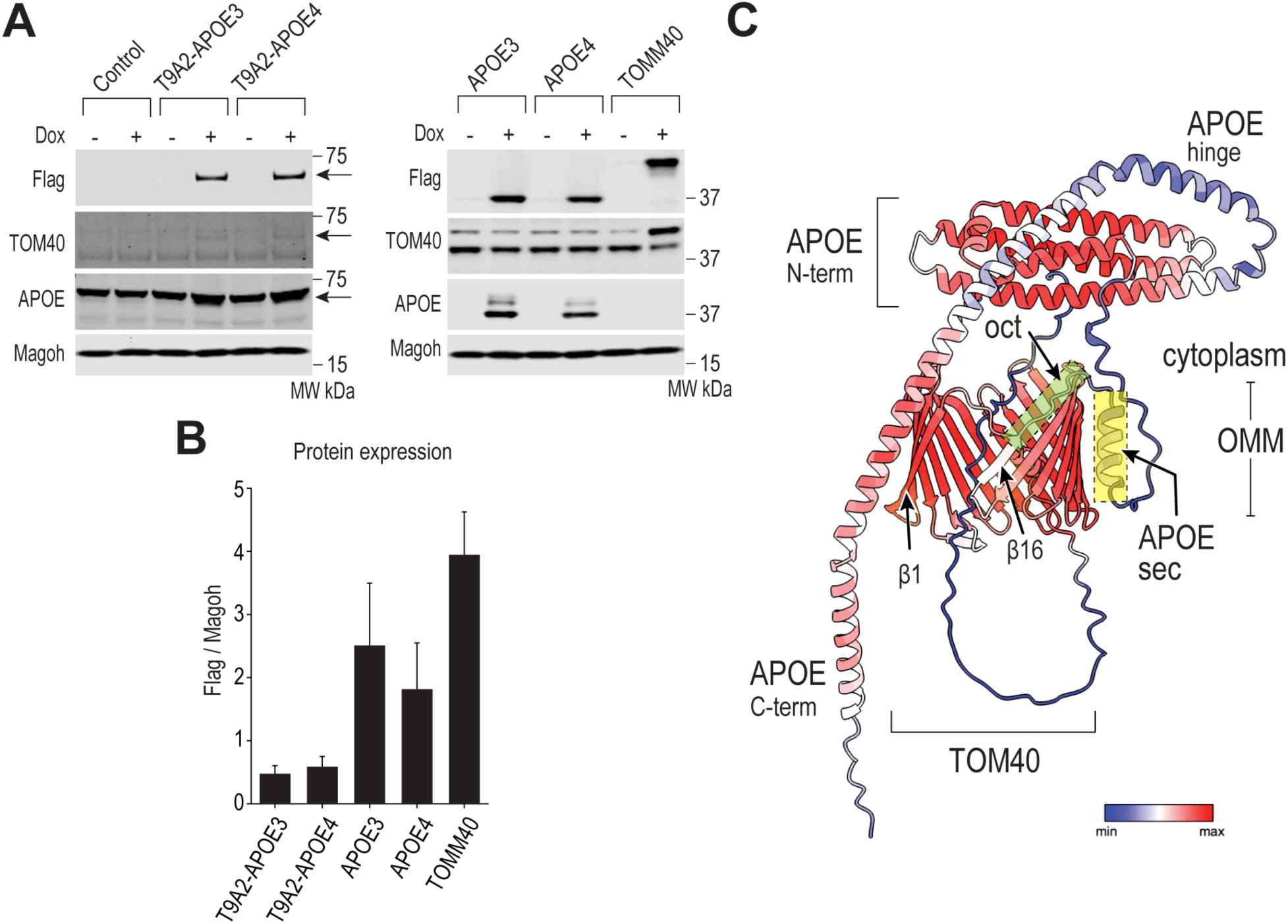
T9A2 mRNA encodes a TOM40-APOE chimeric protein. (**A**). Whole cell lysate of HeLa cells transfected with lentiviruses expressing C-terminal Flag-tagged cDNAs of T9A2-APOE3, T9A2-APOE4, APOE3, APOE4 and TOMM40 under control of doxycycline (DOX)-inducible promoter. Whole cell lysates are prepared at 18h post-induction with 0.5μg/ml DOX. The blots from the same lysates are probed with anti-Flag, anti-TOM40 or anti-APOE antibodies, as indicated. T9A2-APOE3 and T9A2-APOE4 bands (∼70kDa) are indicated by arrow. (**B**) Quantification of expressed protein levels using the anti-Flag and anti-Magoh signals from western blots of the whole cell lysates. Data are presented as mean ± SD, n=3. (**C**) AlphaFold2 predicted T9A2 structural model. Overall view of T9A2 AlphaFold2 predicted model rendered in ChimeraX and colorized by AlphaFold2 confidence metric (pLDDT), min = 19.3, max = 94.4, per-residue on the ribbon structure, as shown in deep red and blue, respectively. The octapeptide (oct) and APOE secretory signal sequence (sec) are indicated in green and yellow boxes with dashed outlines, respectively, and marked with arrows. The TOM40 moiety, TOM40ΔC46, forms a gapped β-barrel lacking β-strands17-19. β1 and 16 are indicated in arrows. TOM40ΔC46 is depicted according to TOM40 experimental structures as immersed in and traversing the outer mitochondrial membrane (OMM) with the apical opening of the transport channel facing the cytoplasm and the bottom face opening to the inner mitochondrial space.

### Structure predictions suggest properly folded APOE tethered to a gapped TOM40 β-barrel

The recently determined cryo-EM structures of *S. cerevisiae* and human TOM40 showed a highly conserved transmembrane beta-barrel channel containing 19 β-strands in which β1 and β19 interact (*50*, *51*). T10 skipping (TOM40ΔC46) deletes part of β16 and β17-19. We used AlphaFold2 (https://alphafold.ebi.ac.uk/), a highly accurate AI-driven 3D protein structure prediction engine (*52*), to generate a template-free structural model of T9A2 with the TOM40ΔC46-oct-sec-APOE3 sequence. In the highest-ranking model (“ranked_0”), TOM40ΔC46 and the N-terminal helical bundle of APOE are modeled in high pLDDT, a per-residue measure of local confidence, while the octapeptide, APOE3 secretory signal and C-terminal domain have low-moderate pLDDT (**Fig. 3C**).

TOM40ΔC46 β1-15 and a half of β16 forms a gapped β-barrel, which superimposes onto human TOM40 (PDB: 7CP9) with C-alpha RMSD 0.853Å. APOE N-terminal helical bundle resembles to the counterpart of the NMR structure of human APOE3 (PDB 2L7B)(*53*) with C-alpha RMSD 1.186Å. The octapeptide, predicted as a coil, takes the place of the missing half of β16. The following APOE secretory signal, predicted as an α-helix, is parallelly close to the β13-15 in the gapped β-barrel, therefore it may be immersed in the OMM, although the model for this region is unreliable. The exact position of APOE, depicted as being in the cytoplasm and interacting with or traversing the OMM, are uncertain, as the tethering sequence oct-seq has lower pLDDT. APOE C-terminal forms a long α-helix of high confidence, but the hinge region has lower confidence. Therefore, the exact position of the C-terminal part relative to the N-helical bundle is plausible, although in the NMR structure the C-terminal helix packed onto the N-helical bundle (PDB 2L7B).

The remarkable similarity of TOM40 and APOE to the experimentally determined atomic structures strongly support the overall structure predictions for T9A2. TOMM40 lacking β17-19 is unlikely to function like normal TOM40, however, it remains to be determined if it could have some function in transport and what interactions with TOM complex and other components it has. Despite having a gapped barrel, T9A2 can be expected to be targeted to mitochondria OMM as N-terminal TOM40 comprising only a few β-strands is sufficient for that as it retains the mitochondrial targeting signals β-hairpins (*54*).

The gap between β-strands 1 and 16 that represents roughly 10-15% of the circumference of the barrel, yet the overall fold and curvature of the β-barrel are nearly identical to that in experimental and AlphaFold2 predicted TOM40 as well as TOM40ΔC46 (**Fig. S2**), suggesting that the β-barrel forms progressively and shaped largely independently of the last closure step effected by interactions of β1 and 19. It is difficult to predict if T9A2 will be properly inserted in the OMM, transport any cargoes, or have other effects on mitochondria, such as by interacting with other TOM components. The predicted folding models of the APOE3 and APOE4 variants in T9A2 are indistinguishable from each other and the same as those of APOE alone pre-calculated in the AlphaFold2 database. However, we cannot rule out that the T9A2-APOE variants have significantly different structures and properties.

### T9A2s associate with mitochondria

We used immunofluorescence confocal microscopy to determine the subcellular localization of the DOX-inducible Flag-tagged T9A2 variants and the corresponding APOEs, detected with anti-Flag antibodies (green channel) (**Fig. 4A**). Both T9A2-APOE3 and T9A2-APOE4 showed similar-sized round bodies throughout the cytoplasm (mean feret diameter of 480-520nm; **Fig. 4B**). The T9A2 staining depended on DOX and there was no T9A2 staining in the nucleus. Importantly, the majority of the T9A2 bodies overlapped with mitochondria, visualized with MitoTracker (red channel), evident in extensive yellow-orange staining throughout confocal optical slices. APOE3 and APOE4 also showed round bodies (**Fig. 4A**), however, they were about 25% smaller than T9A2s and unlike them, had little overlap with mitochondria, evident in fewer yellow areas and 3-fold lower overlap indices (**Fig. 4B**). These observations indicate that T9A2s associate with mitochondria directly, in contrast with APOEs, which are known to take the secretory pathway, involving the ER, Golgi, and transport vesicles. The T9A2s’ subcellular localization is consistent with translation in the cytoplasm and their association with mitochondria is explained by mitochondrial targeting signals harbored in the near-full-length TOM40 (*55*).

**Fig. 4.**
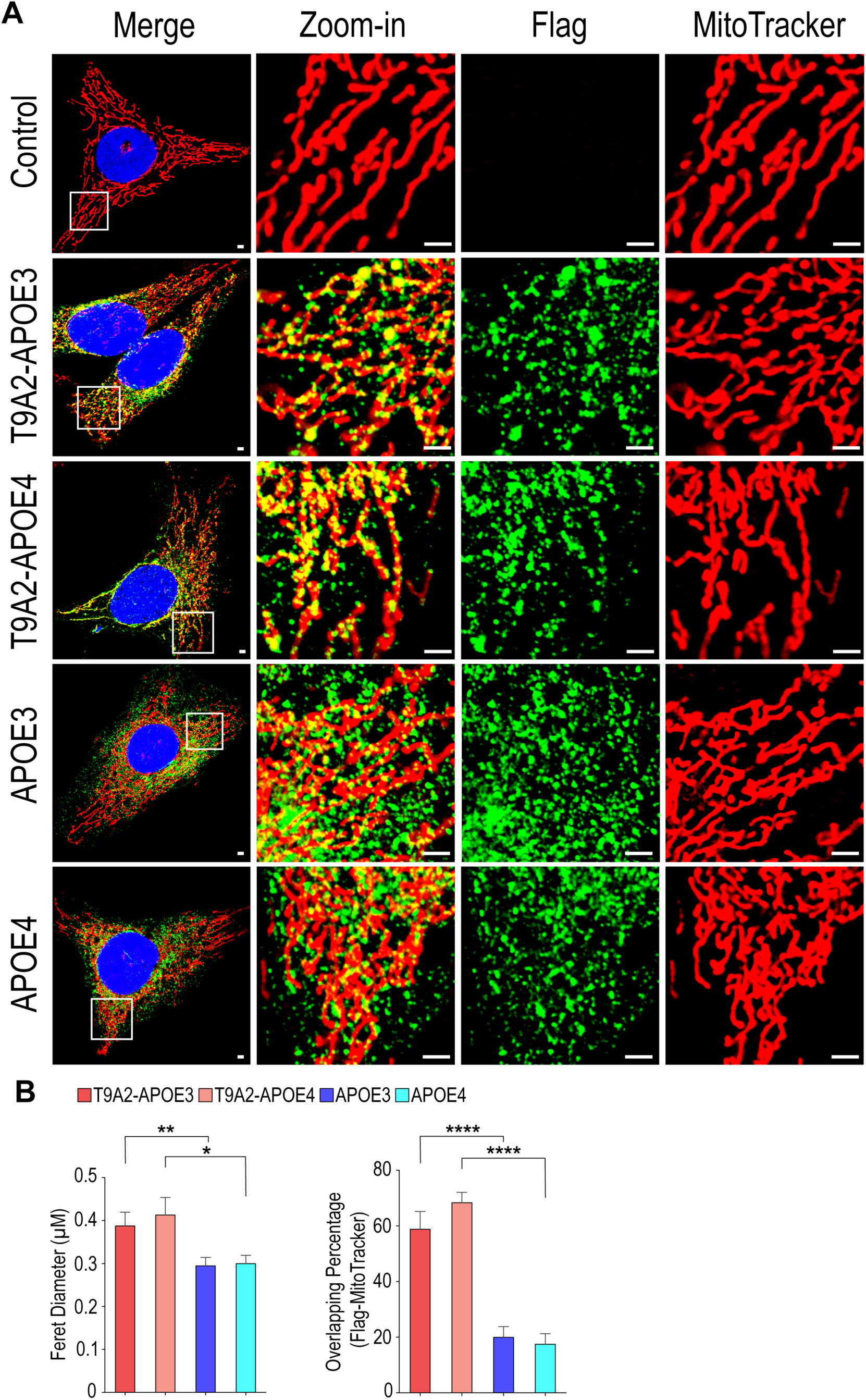
T9A2 associates with mitochondria. Representative immunofluorescence confocal microscopy images (IF) of HeLa CCL2 cells stably transduced with DOX-inducible Flag-tagged T9A2-APOE variants (3 and 4) and the corresponding APOE variants in lentiviral vectors, as indicated, at 18h post-DOX induction (0.5μg/ml). (**A**) Anti-Flag is shown in green and mitochondria are detected with MitoTracker in red. Merge, indicating co-localization in the same optical sections, are yellow-orange. Many of the T9A2 bodies co-localize with mitochondria in adjacent optical sections. Zoom-in views from the T9A2-transfected cells as indicated. 18-25 cells were quantified on 10-12 images per sample. Scale bar, 2μm. (**B**) Histograms showing the T9A2s and APOEs particle sizes (ferret diameter of anti-Flag signal), level of colocalization (percentage of anti-Flag signal with more than 20% pixels overlapping with MitoTracker). Data are represented as means ± SD. n=18-25 cells were quantified on 10-12 images per sample as shown in **A**. One-way ANOVA analyses were performed for the p-value. (*P < 0.05, **P < 0.01, ***P < 0.001 and ****P < 0.001)

### Differential effects of T9A2 and APOE variants on mitochondrial bioenergetic performance

Prompted by the co-localization of T9A2s with mitochondria we investigated if the T9A2 variants affect key mitochondrial bioenergetic functions using the Agilent Seahorse metabolic flux analyzer to determine ATP production rates from mitochondrial respiration (oxygen consumption rate; OCR). Measurements were performed on HeLa cells expressing the above-described C-terminal Flag-tagged T9A2s and APOEs and empty vector as control (**Fig. 5**). T9A2-APOE3, T9A2-APOE4 and APOE3 all increased the basal and ATP production rates by about 2.5-fold and maximal respiration by 3.5-fold compared to control and higher than APOE4, which was only slightly higher than control (**Fig. 5A and B**). Notably, T9A2-APOE3 greatly enhanced spare respiratory capacity (SRC) by 8.7-fold compared with control and higher than T9A2-APOE4, APOE3 and APOE4 (**Fig. 5E**). As previously described, APOE4 SRC was lower than APOE3 (*22*).

**Fig. 5.**
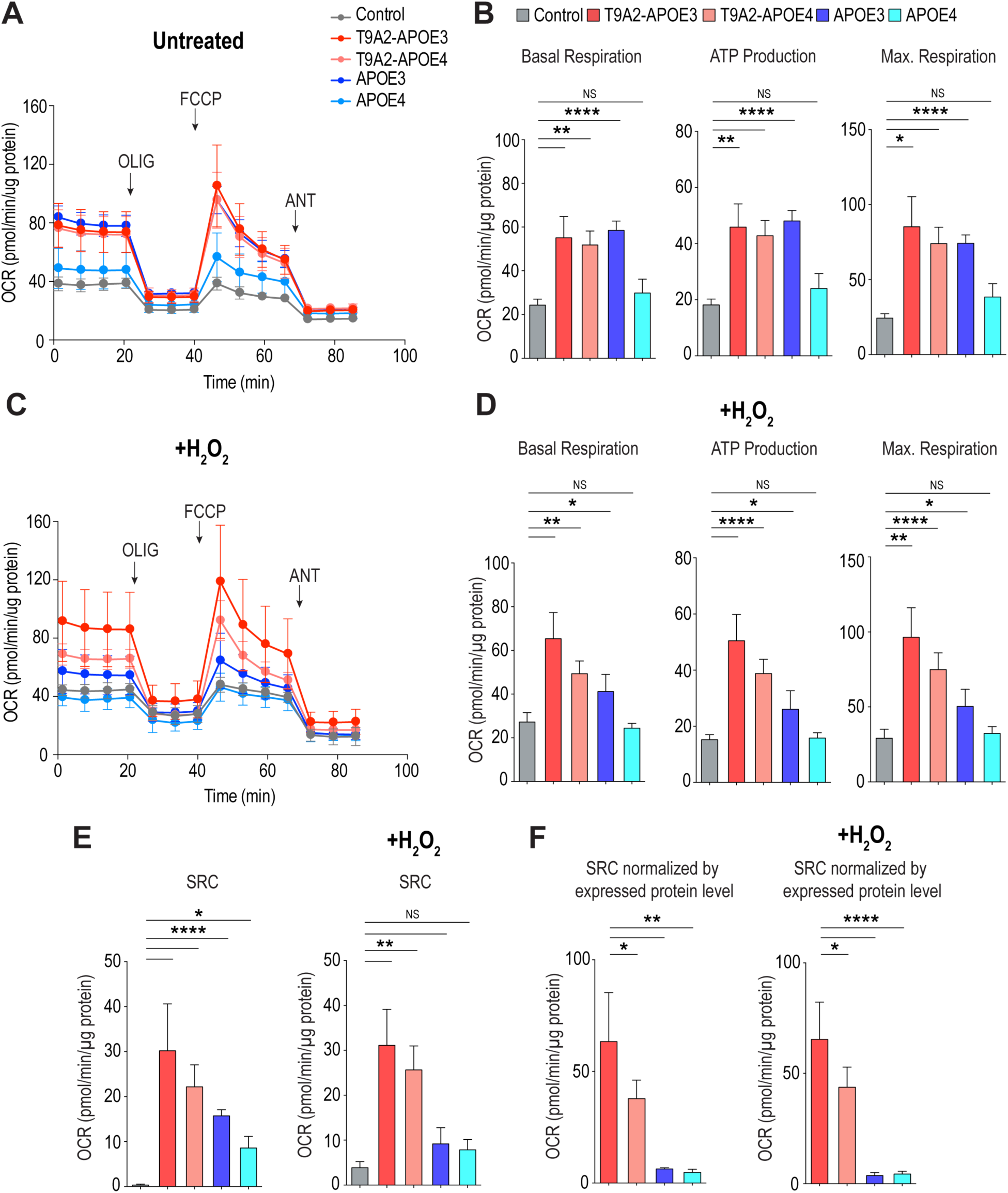
Measurements of T9A2s and APOEs activities on mitochondrial energy production. (**A and C**) Representative graphs of bioenergetic profiles determined in Agilent Seahorse XF analyzer of HeLa cells expressing T9A2-APOE variants (3 and 4) and the corresponding APOE variants, and treated with 100μM H_2_O_2_ for 1h prior to measurements to induce oxidative stress. (**B and D**) Basal respiration, ATP production rate, and maximal respiration (max. respiration) were determined using a mitochondria stress test kit with sequential injections of oligomycin A (OLIG), carbonyl cyanide-p-trifluoromethoxyphenylhydrazone (FCCP), and rotenone+antimycin A (ANT) at the indicated timepoints (arrows) and normalized to the total amount of protein in each well. Biological replicates were performed in parallel on the same number of cells confirmed for each experiment to be in the linear readout range of the tests. (**E**) Spare respiratory capacity (SRC) was calculated as the difference between the maximal and basal respiration. (**F**) SRC values normalized to the corresponding expressed protein levels determined from western blots in Fig. 3B. For all experiments, the bars represent mean ± SD. NS, not significant. One-way ANOVA analyses were performed for the p-value. (*P < 0.05, **P < 0.01, ***P < 0.001 and ****P < 0.001).

SRC reflects the reserve capacity to rapidly increase energy production, which is critically important when there is high energy demand, such as during neuronal stimulation and oxidative stress resulting from reactive oxygen species (ROS) generated by mitochondrial OXPHOS (*56*). To test the effect of oxidative stress we determined the same parameters after treatment with hydrogen peroxide (H_2_O_2_; 100μM for 1h). Under these conditions, basal and maximal respiration, and ATP production rates slightly increased in T9A2-APOE3 compared with untreated cells but decreased significantly in APOE3>APOE4 and remained the same in T9A2-APOE4 (**Fig. 5C and D**). SRC slightly increased with H_2_O_2_ compared with untreated T9A2-APOE3 and T9A2-APOE4 expressers, however, H2O2 treatment led to marked SRC decreases (41.7% and 18.0%) in APOE3 and APOE4, respectively. (**Fig. 5D**). In these measurements, the OCR readings were normalized to the total protein amount in each sample, a common standard. However, as the expression levels of T9A2s were much lower than APOEs (e.g., **Fig. 3**), the SRC fold changes normalized to expressed protein levels were remarkably higher for T9A2-APOE3 versus APOE3 (9-fold and 17-fold in normal and oxidative stress conditions, respectively) and >8-fold higher in both conditions for T9A2-APOE4 compared with APOE3, and APOE4 (**Fig. 5F**). Moreover, normalized basal respiration and ATP production of T9A2-APOE3 were enhanced 4-fold and 1-fold compared with APOE3 (**Fig. S3**), confirming enhanced mitochondria activity and oxidative energy production in T9A2-APOE3, and potentially much more potent than APOE3. These findings indicate that T9A2-APOE3 has biological function that can support mitochondrial health and boost bioenergetic activities, more strongly than T9A2-APOE4, suggesting LOF compared with T9A2-APOE3, and both T9A2s were more potent than APOE3.

### T9A2-APOE3 has anti-oxidant activity that is impaired in T9A2-APOE4

Oxidative stress induces formation of lipid droplets (LDs) which have central functions in lipid metabolism and sequester harmful peroxidized lipids formed by ROS (*57*). APOE3, but not APOE4, has been shown to bind lipids secreted from neurons during oxidative stress and transfer them as apolipoproteins to astrocytes where peroxidized lipids can be detoxified (*58*). We used BODIPY 493/503, a fluorescent lipid staining dye to monitor LDs under oxidative stress (H_2_O_2_ induced). LD accumulation (total area) in T9A2-APOE3, T9A2-APOE4, and APOE3 were comparable under normal conditions but significantly higher in APOE4, nearly 1.5-fold compared with APOE3 (**Fig. S4A and C**), consistent with the previous study that APOE4 induce more lipid droplets accumulation under basal conditions (*13*). With H_2_O_2_, LD accumulation in control and T9A2-APOE3 maintained the same as untreated conditions, and >50% decreased in T9A2-APOE4, APOE3 and APOE4. Double staining of the Flag-tagged proteins and LDs showed that T9A2-APOE3 and T9A2-APOE4 extensively colocalized with LDs in the cytoplasm, as evidenced by overlaps with BODIPY 493/503, showing 2.5-fold higher in T9A2-APOE3 and 1.5-fold higher in T9A2-APOE4 compared with APOE3, and APOE4 (**Fig. S4B and D**). Lipid droplet-associated mitochondria exhibit reduced β-oxidation and display increased ATP synthesis (*59*). This potential interaction of T9A2s with mitochondria and LDs might allow the direct exchange of metabolites and enzymes to produce energy, suggesting a greater capacity of T9A2-APOE3 to sequester toxic peroxidized free fatty acids in LDs.

Since LDs accumulation is altered by increased ROS level, we explored whether accumulated lipids are oxidized. Lipid can be peroxidized in presence of ROS and mediate cellular stress (*60*). To visualize peroxidized lipids, we used BODIPY 581/591 C11, a ratiometric lipid peroxidation sensor, to evaluate changes in lipid ROS. The 510/591nm signal ratio, which indicates the level of lipid peroxidation, was significantly higher in T9A2-APOE4 under normal conditions, suggesting a relatively higher abundance of endogenous peroxidized lipids compared with other constructs and potentially increased sensitivity to fatty acid lipotoxicity (**Fig. S4B**). Upon H_2_O_2_ treatment, the oxidized lipid ratio is slightly increased in control, APOE3, and APOE4. Importantly, the oxidized lipid ratio was significantly decreased in T9A2-APOE3 and less decreased in T9A2-APOE4, resulting in a 33% lower lipid peroxidation level in T9A2-APOE3 compared with control under oxidative stress, while T9A2-APOE4 remains as high as control and APOE3 (**Fig. 6A and 6C**). These data suggest a strong and effective role of T9A2-APOE3 in eliminating H_2_O_2_-induced lipid peroxidation, while the decreased capacity to accumulate LDs and increased lipid peroxidation level in T9A2-APOE4 may result in the LOF for boosting mitochondria respiration during oxidative stress as shown in **Fig. 5F** shows.

**Fig. 6.**
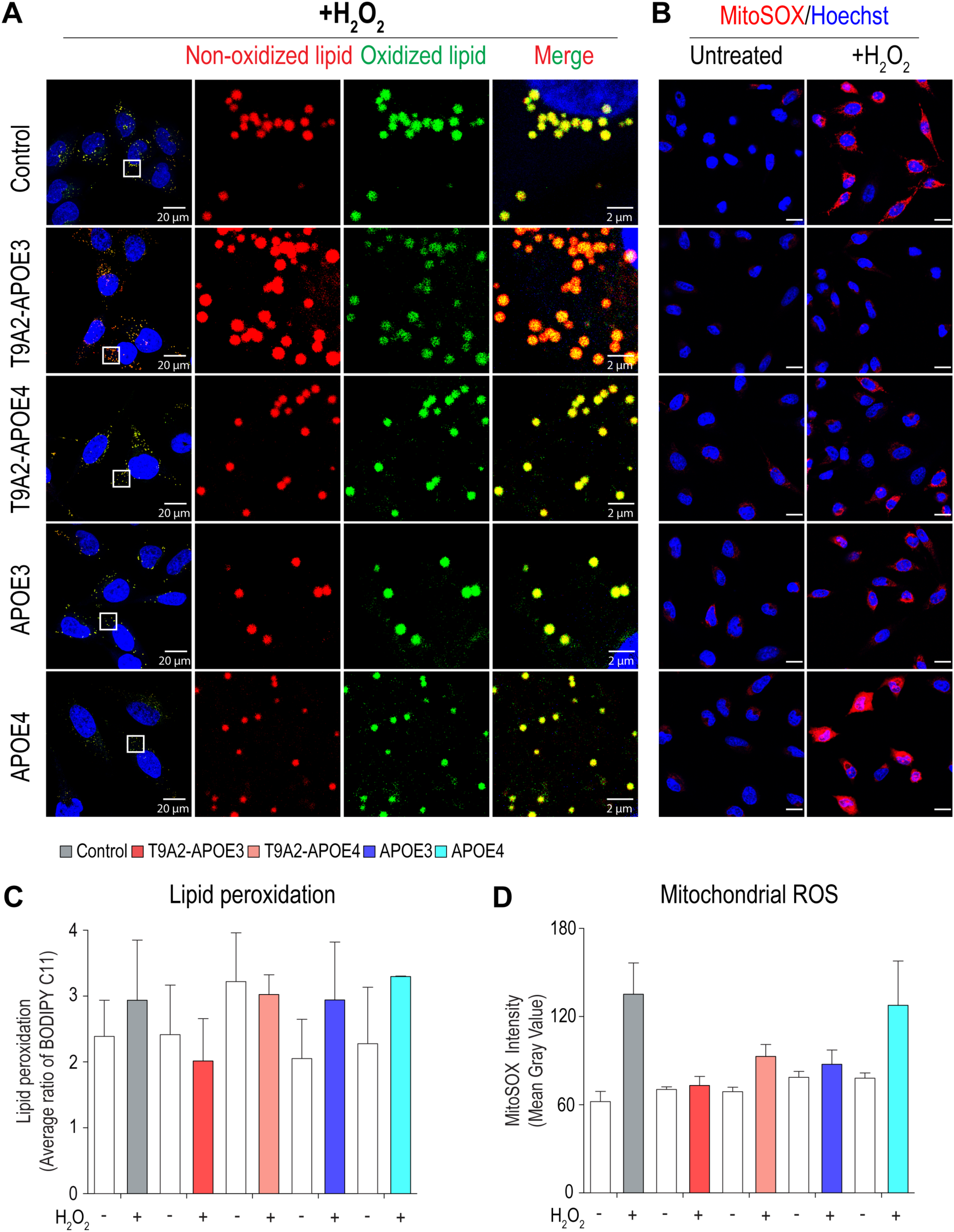
T9A2-APOE3 has higher antioxidant activity compared with T9A2-APOE4 and APOEs. (**A**) Representative immunofluorescence confocal microscopy images (IF) of HeLa CCL2 cells stably transduced with DOX-inducible Flag-tagged T9A2-APOE variants (3 and 4) and the corresponding APOE variants in lentiviral vectors, as indicated, at 18h post-DOX induction (0.5μg/ml), following with 100μM H_2_O_2_ treatment for 1h to induce oxidative stress. The cells were stained with BODIPY 581/591 C11 for evaluation of lipid peroxidation. Non-oxidized lipid (red) and oxidized lipid (green) are shown as indicated. Scale bar, 20μm and 2μm. (**B**) Live cell imaging of HeLa cells expressing T9A2-APOE variants (3 and 4) and corresponding APOE variants incubated with superoxide-sensing MitoSOX after treatment with or without 100μM H_2_O_2_ for 1h. Scale bar, 20μm (**C**) Quantification of lipid peroxidation level, expressed as the average ratio of the intensities of the 510 (green) and 591 nm (red) emissions quantified from the images in **A** and **Fig. S4B**. n=12-15. (**D**) Quantification of MitoSOX fluorescence intensities (mitochondrial ROS level) from the images in **B**. n=12-15. For all experiments, the bars represent mean ± SD. One-way ANOVA analyses were performed for the p-value. (*P < 0.05, **P < 0.01, ***P < 0.001 and ****P < 0.001).

Mitochondrial ROS production was monitored with MitoSOX, a superoxide-sensitive fluorogenic dye specifically targeted to mitochondria in live cells. As shown in **Fig. 6B**, T9A2-APOE3-expressing cells had the lowest MitoSOX staining compared with untreated cells, T9A2-APOE4, and the APOEs (low to high MitoSOX ranking T9A2-APOE3<T9A2-APOE4/APOE3<APOE4/control) in cells treated with H_2_O_2_. The production of mitochondrial superoxide in T9A2-APOE3 were 55% and 15% lower than control and APOE3, respectively, and T9A2-APOE4 was 40% and 26% lower than control and APOE4, respectively (**Fig. 6D**). Thus, T9A2-APOE3 has anti-oxidant activity that can protect mitochondria from excessive ROS and dysfunction more effectively than T9A2-APOE4 and the APOEs.

### The interactomes of T9A2s and APOEs may explain APOE4 and T9A2-APOE4 roles in mitochondria dysfunction and neurodegenerative diseases

We performed immunoprecipitations (IPs) with anti-Flag from the cell lines expressing the Flag-tagged APOEs, T9A2s, and TOMM40 (as reference), and empty vector (background control) to profile their protein interactomes by label-free quantitative mass spectrometry (MS). The fold change (FC) of significantly enriched proteins (|log2FC|>1.0; p-value<0.05) averaged from biological replicates of each are listed in supplementary table **S9**. In each sample the IP target(s) was the most abundant component, APOE and/or TOMM40. The MS did not distinguish between the APOE variants. Venn diagrams showed more target specific differences than commonalities in pairwise and multiple comparisons of co-IP proteins (**Fig. S5**).

The significantly enriched proteins from each target were analyzed and visualized using STRING database of known and predicted protein–protein interactions, including STRINGdb’s highest confidence predicted functional partners (<6, indicated in each co-IP list in **Data S9**). The complete networks for each target sorted into clusters are shown in **Fig**. **7**. Clusters containing the target protein (APOE and/or TOMM40) and strongest interactors are shown in **Fig. S6.** All the co-IP components of each cluster were further analyzed for gene ontology, functions and pathways (GO, KEGG, Reactome, and WikiPathway) and the results are displayed in **Fig. S6.** The datasets are rich in information and consistent with many known interactions (indicated by the connecting lines that link to STRINGdb’s evidence sources) and functions of TOMM40 and APOE, validating the accuracy of our methodology. Several themes are noted here.

**Fig. 7.**
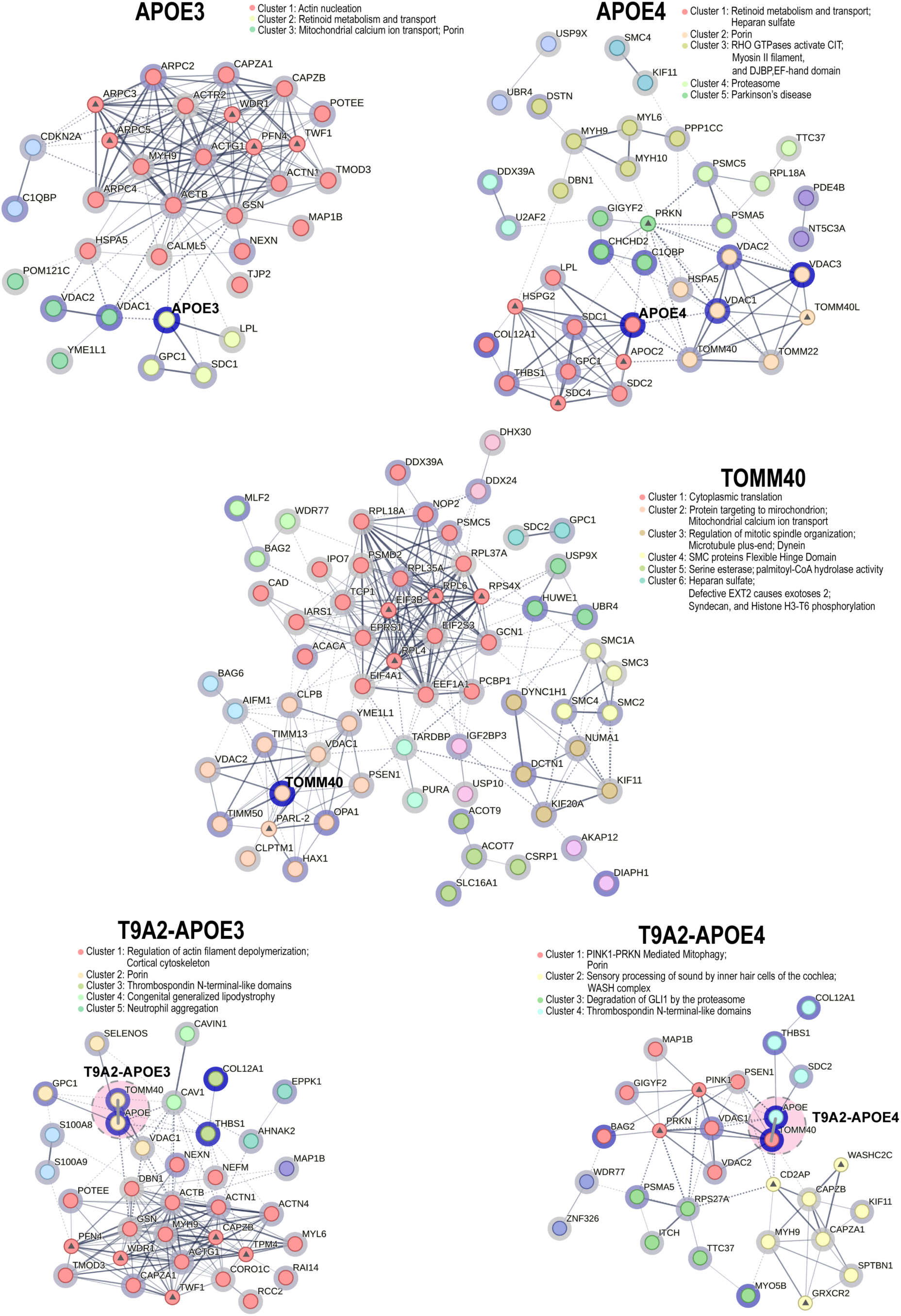
T9A2s protein interactomes. STRING network analysis of IP-MS significantly enriched proteins (log2FC>1, p-value<0.05) identified in APOE3, APOE4, T9A2-APOE3, T9A2-APOE4 and TOMM40. Each protein is represented by its gene name as a round node, which is encircled in blue of intensity corresponding to their abundance (log2FC enrichment rank) in that IP. The nodes with a central triangle represent predicted functional partners from the STRING database. APOE and TOMM40 in T9A2-APOE3 and T9A2-APOE4 are shown as touching with their centers connected by a short thick line to indicate they are in fact one protein and highlighted in an ellipsoid. The interacting nodes are connected by edges which indicates both physical and functional protein-protein interactions from the STRING database, where edge thickness correlates with interaction score with a medium confidence set at 0.4. The functional clustering was performed using the MCL clustering algorithm in the STRING databases and the classification are shown alongside the network. Subclusters of the network are colored nodes as indicated.

As expected, TOMM40 co-IP many mitochondrial components, including the OMM β-barrel proteins, VDAC1 and VDAC2 (Voltage-dependent anion-selective channel proteins), the IMM translocase complex’s TIMM50 and TIMM13, and OPA1, YME1L1, and CLPB, which control mitochondria morphology, dynamics, and fate, and HAX1 an apoptosis regulator. It is uncertain if the cluster of cytoplasmic ribosomal proteins and translational factors co-IP with TOMM40 represent specific interactors considering TOMM40’s several-fold higher expression level than the other targets. Consistent with APOE’s functions, both APOE3 and APOE4 highly enriched for retinoid metabolism and transport cluster, highlighted by lipoprotein lipase (LPL), and syndecan (SDC1) and glypican (GPC1), cell surface heparan sulfate proteoglycans (HSPGs) that link the cytoskeleton to the interstitial matrix. SDC1 is also found in lysosomes and GPC1 is also associates with CAV1 (caveolin1)-containing vesicles. APOE4 had more extensive interactions with the same group, including SDC2, SDC4, and HSPG2/perlecan, and interactor THBS1, all of which have roles in focal adhesions, motility and axonal regenerations. The T9A2s do not interact with LPL or enrich for the retinoid metabolism and transport cluster.

Most of the mitochondrial proteins enriched in TOMM40 IP were not enriched in the IPs of T9A2s or APOEs, suggesting that they do not embed or interact with mitochondria the same as TOMM40. VDACs were detected in all the targets and particularly in APOEs where they were among the top co-IP proteins and several-fold more enriched (FC) and higher ranked in the co-IPs of the T9A2s and TOM40 IPs. The best characterized, VDAC1, is a nexus point for cell death pathways and a key mediator of energy metabolism, including ATP/ADP exchange between mitochondria and the cytoplasm. It is also a part of IP3R-GRP75-VDAC1 complex that bridges the mitochondria-associated ER membrane (MAM) to mitochondria, which regulates Ca2+ transfer between them and thereby activity of mitochondria (*61*). The MAM could potentially facilitate access of APOEs from the ER to mitochondria. Strikingly, APOE4 has more extensive interactions with mitochondria than APOE3, including with the TOM complex’s TOMM40 and TOMM22, the cytoplasmic receptor for pre-mitochondrial sequences, consistent with APOE4 routing through the cytoplasm. Additionally, APOE4 co-IP CHCHD2, an IMM protein and stress-induced regulator of transcription of cytochrome c oxidase 4I2, a subunit of the terminal enzyme of the electron transport chain, and regulator of mitochondria-mediated apoptosis. APOE4 and T9A2-APOE4 were also enriched in ubiquitin-proteasome components.

Multiple differences between T9A2-APOE3 versus APOE3 or T9A2-APOE4 versus APOE4 involved cytoskeletal components of actin filaments, microtubules and motor proteins, suggesting differential effects on intracellular transport of organelles (including mitochondria and endosomes) and various macromolecular complexes (**Fig. 7**). T9A2-APOE3 but not T9A2-APOE4 also enriched for RHOB GTPase cycle, which plays an important role in neural connectivity and cognition and is involved in AD pathogenesis. This is consistent with a previous report that APOE3 exclusively upregulates the focal adhesion and actin cytoskeleton pathway (*62*) while APOE4 showed actin cytoskeleton dysfunction and decreased endocytosis (*63*).

Pathway analysis noted high enrichment in Porin activity (OMM’s TOMMs and VDACs) across the targets, and PINK1-PRKN-mediated mitophagy, a hallmark Parkinson’s disease pathway, especially in T9A2-APOE4 and to a lesser degree in APOE4 (**Fig. S6**). This pathway involves Parkin (PRKN), the PINK1-controlled mitophagy-promoting E3 ubiquitin ligase that ubiquitinates OMM proteins, is also linked to AD (*64*). T9A2-APOE4 also enriched for ALS, and negative regulation of oxidative stress. T9A2-APOE3 also enriched for congenital generalized lipodystrophy and cholesterol metabolism. TOMM40 enrichment in mitochondrial organization and transport, and mitochondrial functions, as expected.

## Discussion

The fusion of two of the highest risk genes in age-related cognitive decline, AD, and longevity, APOE and TOMM40, is an unforeseen scenario with potentially profound implications for cell physiology, cellular defense, and multiple diseases. Any regulation of RNAs produced from the TOMM40-APOE genomic region is therefore of great interest. T9A2 may have escaped detection (or failed to receive deserved attention if it had been) due to its low abundance and possibly transient expression. However, low abundance does not deny biological significance; in fact, it may be a feature of late onset diseases and may explain their exceedingly slow progression, requiring decades to manifest. A pathogenic process that slowly eliminates neurons, a minimally replenishable cell population, may only reach a tipping point after a very long time, after most of the affected neurons have been eliminated.

Numerous studies have implicated APOE4 and TOMM40 in mitochondria dysfunction, which precedes by decades neurodegeneration, dementia, and accumulation of amyloid plaques and neurofibrillary tangles. Yet, massive research and development costs have been aimed at preventing or clearing these late disease manifestations that have not yielded effective therapies. Controversies concerning the role of TOMM40 and a lack of understanding how APOE4 secreted protein, impairs mitochondria have hampered progress towards development of therapies based on these components. The discovery of chimeric TOMM40-APOE, T9A2, and its functions could open up new opportunities for advancing basic understanding of AD mechanisms and prospects of new therapies.

T9A2s add new actors to the APOE sphere and may explain why it has been so challenging to explain the bewildering myriad of triggers and diversity of molecular phenomena in AD and other diseases impacted by the TOMM40-APOE locus. The multiple differences in properties and interactions of T9A2-APOE3 and T9A2-APOE4 and the corresponding APOEs and TOMM40, have the potential to create a great complexity of scenarios depending on APOE variants and TOMM40 polymorphisms, and could vary in different cell types and conditions. The ability of T9A2-APOE3 to boost SRC and dampen oxidative stress, are highly beneficial and exceed those of APOE3. This suggests a rationale for the readthrough-splicing as a strategy to enhance cell resilience and ability to survive adverse conditions. T9A2-APOE4 is inferior to T9A2-APOE3 in this respect, however, it may in some measures be advantageous to APOE. T9A2-APOE4 is an alternative source and direct route of APOE4 to mitochondria. T9A2-APOE4 and its interactome enriches for Parkinson’s disease associated PINK1-PRKN mediated mitophagy pathway. This and the rest of the interactomes’ data have a wealth of leads for further investigation.

Importantly, APOE3 and T9A2-APOE3 could work in concert and possibly synergistically, to alleviate oxidative stress by expelling peroxidized lipids and sustain mitochondrial health, respectively. The lack of APOE3 in homozygous APOE4 or its decrease in heterozygotes creates a vulnerability, especially in high energy demanding neurons from oxidative stress, lower ability to boost energy production, and expel harmful peroxidized lipids, would make high energy demanding cells, like neurons, highly vulnerable and could cause injury and death. Exogenous APOE4 could exacerbate this further.

A schematic of gene expression from the TOMM40-APOE genomic region and their biological outcomes is shown in **Fig. 8**. T9A2 and APOE could both be induced by cell stress due to read-through from TOMM40 weak terminal PAS and de novo APOE transcription from its own promoter upstream of exon1, as a concerted counter-measures to oxidative stress. APOE3 could ameliorate the effect of excessive lipid peroxidation by facilitating transfer of lipids from neurons through the secretory pathway to glia cells that can detoxify them, while T9A2-APOE3 could reduce ROS and boost mitochondrial bioenergetic capacity that is needed under the same conditions. Both activities are impaired by the APOE4 variant, and their combined LOF could create a particularly harmful environment in neurons. Importantly, T9A2s and APOEs act at different locations and T9A2-APOE3 stronger beneficial effects compared with APOE3 suggest a rationale for forming T9A2 chimera to enhance cell survival during acute oxidative stress. It is uncertain if T9A2-APOE4 is only a LOF relative to T9A2-APOE3 or also GOF toxicity with respect to other functions that we have not investigated here. It will be interesting to know how T9A2s exert their functions at the OMM, including whether the T9A2 gapped β-barrel has any transport activity itself and whether it affects other TOM subunits or OMM constituents.

**Fig. 8.**
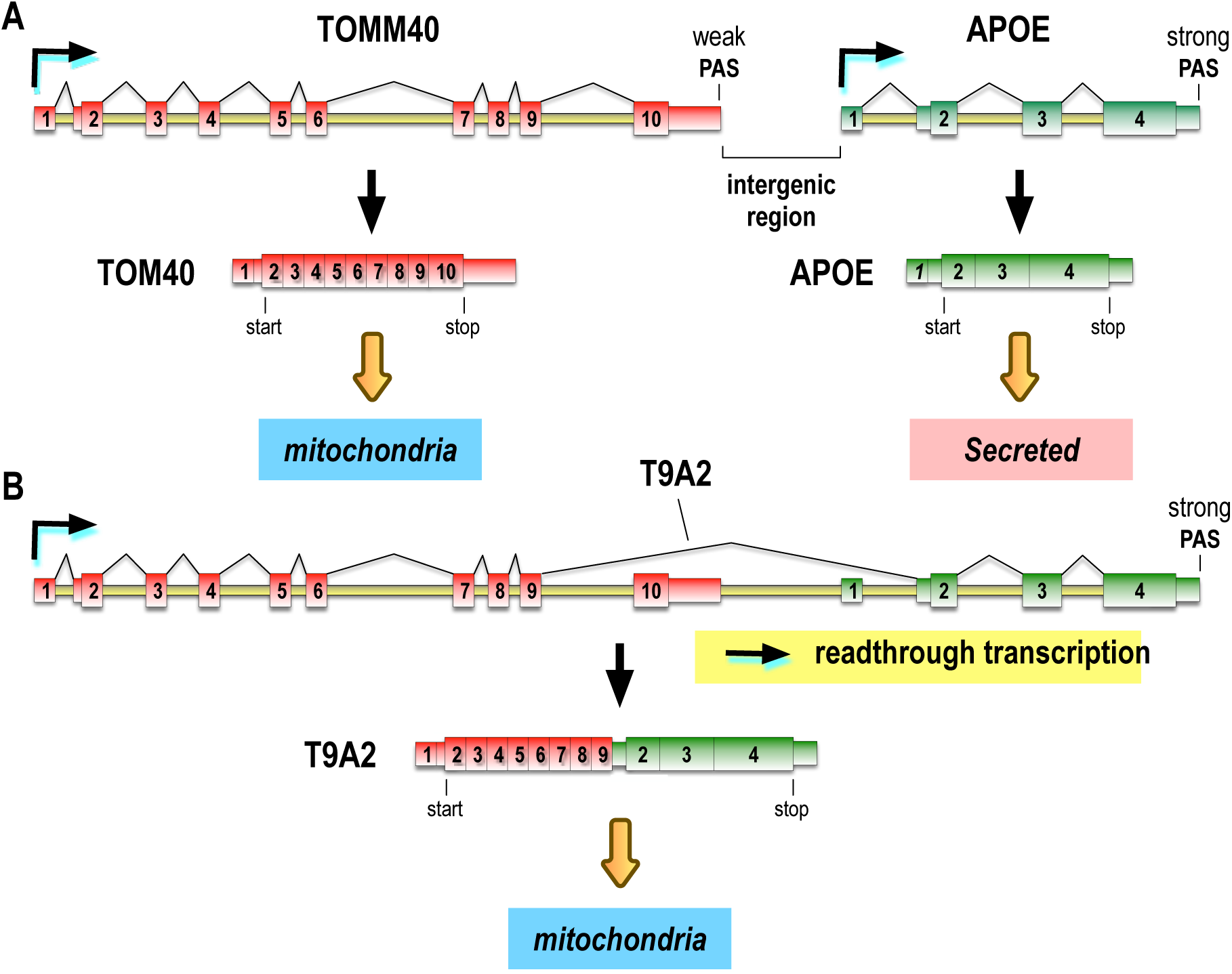
A schematic diagram of the mRNAs and proteins that can be produced from the TOMM40 and APOE genes. (**A**). TOMM40 is essential for mitochondria and cell viability and transcriptionally active in all cells. APOE is only expressed in some cells but can become transcriptionally active under cell stress from its own promoter in other cells, like neurons and HeLa, and produce a secreted protein. (**B)** Cell stress also causes readthrough transcription from TOMM40 that extends through the downstream intergenic region to the end of APOE, yielding TOMM40-APOE pre-mRNA that can splice its introns, including chimeric splicing, primarily TOM40Δex10-APOEΔex1, called T9A2 that makes a fusion protein, TOM40ΔC46-oct-sec-APOE in which full-length APOE is in-frame with near full-length TOM40. The fusion protein includes an octapeptide (oct) translated from the 5’-UTR in APOE exon2 and the APOE secretory signal sequence (sec), which is otherwise cleaved from APOE that is transcribed from its own promoter as it is translated into the ER for secretion. However, T9A2 is translated in the cytoplasm and targeted to mitochondria, providing a direct pathway for delivering APOE variants to mitochondria. TOMM40 readthrough alone can down-regulate TOM40 expression. The bent arrows with turquoise shading indicate transcription activity and direction. Splicing patterns and resulting mRNAs are indicated with blue arrows. Square arcs connecting splice sites indicate splicing events.

Chimeric cis-splicing in readthrough transcripts that span gene pairs have been described in many genes sometimes referred to as conjoined genes (*65–67*). Only a fraction of these have potential ORFs, and only few make detectable proteins, usually at very low levels, which have made them challenging to study. While it is intriguing that T9A2 mRNAs appear to be up-regulated under stress conditions and viral infections, and influenced by genetic backgrounds, they come from small studies and will need to be tested in much larger cohorts.

T9A2 also appears to address the long-standing controversy about TOMM40 role in AD, stemming from GWAS in which APOE4 alone, without pathogenic mutations in TOMM40, was sufficient to explain AD risk. However, T9A2, which was not known at the time, suggests an alternative and new role for TOMM40 as a driver of APOE4 expression as a fusion protein. We propose that this principle could be a general post-transcriptional pathogenesis mechanism whereby readthrough transcription from a normal gene activates a downstream risk gene that may otherwise not be expressed in the same cells.

While this manuscript was in final preparation, we learned about a report from Chang et al (*68*) described identical TOMM40-APOE chimera from deep RNA-seq and showed their localization to mitochondria. However, Chang et al. have not detected differences between T9A2-APOE variants’ effects on mitochondrial functions or relate T9A2-APOE4 to disease pathogenesis.

## Methods

### Cell culture, transfection and treatment

Cells (Human cervical carcinoma HeLa, HeLa CCL2, and 293T) were obtained from ATCC and maintained in Dulbecco’s modified (DMEM) supplemented with 10% fetal bovine serum (FBS), 10 unit/ml penicillin and 10µg/ml streptomycin at 37°C and 5% CO_2_. The HeLa CCL2 Tet-inducible cell lines were maintained in regular media supplemented with 10% Tet-system-approved FBS (Takara Bio, 631101) and 1% penicillin–streptomycin. All cell lines were tested for mycoplasma contamination. The sequences of the U1 and control AMOs (Gene Tools) are 5′-GGTATCTCCCCTGCCAGGTAAGTAT-3′ and 5′-CCTCTTACCTCAGTTACAATTTATA-3′, respectively (*69*). The AMOs were transfected into 1×10^6^ cells using a Neon electroporation system (Invitrogen) according to the manufacturer’s instruction to achieve the desired final doses indicated in the text and figures. 80nM THZ531 (Cayman Chemical) CDK12 inhibitor dissolved in DMSO was added to the culture medium for 8-12h.

### Lentiviral expression of chimeric TOM40-APOE

The coding sequences of APOE3, APOE4, TOM40, chimeric TOM40-APOE3 and TOM40-APOE4 fused at their 3’-ends to 3×Flag were synthesized from Genewiz and inserted into pCW57.1 (Addgene #41393) with EcoRI (ThermoFisher Cat. FD0274) and BamHI (ThermoFisher Cat. FD0054). Lentivirus was produced by transient transfection of pCW57-TOM40-APOE3 or T9A2-APOE4 into 293T cells along with the envelope plasmids pMD2.G (Addgene Cat. 12259) and psPAX2 (Addgene Cat. 12260) with Lipofectamine 3000 (ThermoFisher catalog L3000001) according to the manufacturer’s instruction and harvested after 24h transfection. HeLa CCL2 cells were infected by the lentivirus in 6-well plate at 75-80% confluency and continuously selected in the medium containing 1µg/ml puromycin (ThermoFisher catalog A1113802) for 24h. TOM40-APOE expression was induced by growing the culture in present of 0.5µg/mL doxycycline (DOX) for 24h.

### Reverse transcription and PCR amplification of TOMM40-APOE readthrough transcripts

Total RNA was extracted from culture cells using TRIzol reagent (ThermoFischer Scientific 15596026). SuperScript™ IV VILO™ Master Mix (ThermoFischer Cat. 11756050) was used to generate cDNA. 1µg of total RNA was utilized as template in a 20 micro-liter reaction. A pair of primers, T9A2_Fp 5′-CCCGTCCACTTTCCCATCTC-3′ and T9A2_Rp 5′-GCTCCTGGGGAAGGACGTC-3′, and 2% of the cDNA were used in TOMM40-APOE readthrough transcripts amplification PCR reaction. Amplicons were subsequently separated on 1% agarose gel.

### Western blots

Whole cell lysate, sample processing, SDS-PAGE and immunoblotting were performed as previously described (*70*). In detail, 24h post treatment, HeLa cell in adherent was washed briefly in cold PBS, solubilized in 200µl 1×SDS sample buffer for the cell in one well out of a 6-well plate, and passed through a 27-gauge needle 5 times to shear the genomic DNA. Whole cell lysate was heat to 95°C for 10min, separated on NuPAGE 4-12% Bis-Tris gels (Invitrogen), and transferred to nitrocellulose membrane with iBlot2 Dry Blotting System (ThermoFisher). Mouse anti-Flag antibody (Sigma, F1804), Rabbit anti-TOMM40 antibody (Abcam, ab188543), rabbit anti-APOE antibody (Sigma-Aldrich HPA068768), rabbit anti-VDAC1 (Abcam, ab15895), rabbit anti-tubulin (Cell Signaling Technology, 2126S) and mouse anti-Magoh antibody (21B12) (*71*) were used in 1:1000, 1:1000 and 1:500 dilution, respectively. The membranes were scanned on an Odyssey® infrared imaging system (Li-Cor), and the intensity of the protein bands was analyzed using the software provided by the manufacturer.

### RNA-seq

Total RNA extraction from cells was done at either 6 h or 24 h after AMO transfection using Trizol (Invitrogen) followed by either poly(A) RNA purification on oligo-dT beads using Dynabeads™ mRNA Purification kit (ThermoFisher) or ribosomal RNA depletion using the Ribo-Zero kit (Illumina). Nascent RNAs were metabolically labeled in cells with 250µMol 4-thiouridine (4-shU) that was added for 30min at 7.5–8h post AMO transfection. 4-shU-containing mRNAs, the free thiols on poly(A) mRNAs were reacted with 0.2mg/ml of EZ-link biotin-HPDP (Thermo Scientific) for 1h, then purified on M-280 streptavidin Dynabeads (Invitrogen), as previously described(*36*). cDNA synthesis and RNA-seq library preparation were constructed either using the KAPA stranded RNA-seq library preparation kit (Roche) according to the manufacturer’s instructions. The sequencing was performed on Illumina NovaSeq 6000 platforms.

### RNA-seq data analysis

Paired-end RNA-seq reads were trimmed of any adaptor sequences with the FASTX-Toolkit (version 0.0.14). The two paired reads were merged into one single fragment using PEAR (version 0.9.8), and then fragments larger than 150nt were filtered out. The remaining reads were aligned to the GRCh38/hg38 reference genome using STAR (version 2.5.3a) with the following parameters: –twopassMode Basic –alignSJoverhangMin 5 –alignSJDBoverhangMin 5 – outSAMmapqUnique 255 –outFilterMultimapNmax 1 –outSJfilterReads Unique. Reads per exon were grouped, from which RPKM (reads per kilobase per million mapped reads) values were calculated using SAMtools (version 0.1.19). In order to directly compare samples that have a different number of mapped reads, the read coverage for each sample was normalized to the total number of mapped reads per million (RPM). This normalized value was also used to scale the samples for visualization on the UCSC Genome Browser (http://genome.ucsc.edu/). External RNA-seq datasets were downloaded in raw format from the Gene Expression Omnibus (https://www.ncbi.nlm.nih.gov/geo/) and then processed and aligned as described above. Sashimi plots were rendered using ggsashimi (*72*).

### BLAST search in NCBI Sequence Read Archive (SRA)

RNA-seq datasets were first selected by keyword queries in the SRA database at NCBI (https://www.ncbi.nlm.nih.gov) and metadata were obtained using the built in Run Selector option. Blast search in selected SRA dataset was done by blastn_vdb program in NCBI SRA Toolkit (GitHub – ncbi/sra-tools: SRA Tools) on HPC in PMACS with the specified parameters - strand=”plus” -outfmt “6 sacc saccver sseqid slen sstart send qlen qstart qend qcovs evalue bitscore score pident nident mismatch positive sseq” -num_threads 4 -task “blastn” - max_target_seqs 50000 -evalue 10 -sra_mode 2 and the rest parameters in their default. Blastn_vdb was embedded in homemade python scripts to facilitate array job submission onto HPC. After blast search finished, to filter the blast results to get hit, E-value ≤1E-6, pident ≥95% and qcovs≥90% was used for the RNA-seq datasets of reads shorter than or equal to 70% of query length, and E-value ≤1E-6, pident ≥95% and qcovs ≥80% was used for the RNA-seq datasets of reads longer than 70% of query length.

### AlphaFold2 structure prediction

An AlphaFold2 GPU workstation is set up on AWS EC2 by following the tutorial at https://github.com/deepmind/alphafold and AWS EC2 reference. T9A2 is predicted with the customized parameters ‘-max_template_date’=2017-08-01, in order to avoid using determined TOM40 structures as model template, and the FASTA sequence of T9A2, the other parameters are used their default.

### Immunofluorescence (IF) confocal microscopy

IF was performed on HeLa (ATCC clone CCL2) cells transduced with the seeded in glass bottom 6-well plates (2×10^5^ cells/well) incubated in DMEM medium containing 0.5µg/mL DOX for 18h. 1mM MitoTracker diluted 1:2000 ratio in DMEM was added for 10min, following which the cells were washed twice with PBS and fixed with 300uL formaldehyde solution (ThermoFisher, Image-iT™ Fixative Solution) for 15min. The cells were washed twice with PBS and permeabilized with 0.2% Triton X-100 in 300 µl PBS with for 10 min, washed twice with PBS and incubated with 1mL blocking solution (Li-Cor, Intercept PBS Blocking Buffer) for 1.5h followed by two additional PBS washes. Staining was carried out with anti-Flag antibody (Sigma, F1804) 1µg/mL in PBS for 1h, two washes with PBS, and Fluro-488-conjugated secondary antibody (Invitrogen, A-11001) 2µg/mL in PBS for 1h. For BODIPY 493/503 staining, 10µM BODIPY 493/503 (Invitrogen, D3922) was diluted in PBS and incubated for 15min. Following two additional PBS washes the cells were incubated with nuclear staining DAPI solution (NucBlue Fixed Cell Stain ReadyProbes reagent), and sealed with a cover glass. Imaging was performed using Zeiss confocal microscope LSM880 and images were processed identically with Fiji ImageJ software.

### Mitochondrial respiratory activity assay

Mitochondria respiration activity in HeLa cells expressing C-terminal Flag-tagged APOE3, APOE4, TOMM40, T9A2-APOE, T9A2-APOE4 same empty vector as reference was measured by using the Agilent Seahorse XF Cell Mito Stress Test Kit (Agilent Technologies, 103015-100) and Seahorse XFe96 FluxPak (Agilent Technologies, 103793-100), and the oxygen consumption rate was monitored with the Seahorse XFe96 Analyzer (Agilent Technologies) following the manufacturer’s instructions. HeLa cells (6×10^4^/well) of the indicated genotypes were seeded in a 96-well plate and cultured with medium containing 0.5μg/mL DOX on the day before the assay including four to six replicate culture wells per run. One hour before the assay, cells were washed once with Seahorse XF DMEM medium pH 7.4 (Agilent Technologies, 103575-100) followed by addition of 180μl Seahorse XF DMEM medium pH 7.4 per well, supplemented with 1mM pyruvate, 2mM glutamine and 20mM glucose (Agilent Technologies, 103578-100, 103579-100 and 103577-100), cells were cultured for 1h at 37°C in a CO_2_-free incubator. The Mito-Stress test was performed following the standard protocol. Standard mitochondria stress tests were performed by first measuring basal values followed by measurements after sequential addition of 0.8µM oligomycin, 0.8µM FCCP, and 0.5µM rotenone/antimycin A. After the assay, protein concentrations of each well were determined via BCA protein assay (ThermoFisher, 23227) and used to normalize oxygen consumption rate values.

### Live-cell mitochondrial ROS measurements

HeLa (ATCC clone CCL2) cells transduced with the seeded in glass bottom 6 well plates (2×10^5^ cells/well) incubated in DMEM medium containing 0.5µg/mL DOX for 18h. After washing once by PBS, pre-warmed DMEM medium containing 100µM H_2_O_2_ was added to the cells for 1h treatment. After washing twice by HBSS, 5µM MitoSOX Red (Invitrogen, M36008) and 2mg/ml Hoechst 33342 (Invitrogen, H3570) in DMEM were added to cells and incubated for 15min at 37°C. After washing twice by HBSS, FluoroBrite™ DMEM (Thermo-Fisher Scientific, A1896701) supplemented with 10mM HEPES and 4mM glutamine was added to cells, and images were acquired by Zeiss confocal microscope LSM880 in a chamber heated to 37°C in 5% CO_2_. The fluorescence intensities of MitoSOX Red were determined by ImageJ.

### Lipid peroxidation assay

HeLa (ATCC clone CCL2) cells transduced with the seeded in glass bottom 6 well plates (1×10^5^ cells per well) incubated in DMEM medium containing 0.5µg/mL DOX for 18h. After washing once by PBS, pre-warmed DMEM medium containing 100µM H_2_O_2_ was added to the cells for 1h treatment. The cells were then incubated with 10µM BODIPY 581/591 C11(Invitrogen, D3861) at 37 °C for 30 min, followed by fixation with 4% formaldehyde solution (ThermoFisher, Image-iT™ Fixative Solution) for 15min, the nucleus were stained by DAPI solution (NucBlue Fixed Cell Stain ReadyProbes reagent) and the images were acquired by Zeiss confocal microscope LSM880. The fluorescence intensities from both non-oxidized C11 (591nm emission) and oxidized C11 (510nm emission) were measured. The mean fluorescence intensity of the two channels were determined by ImageJ. It should be noted that the BODIPY 581/591 C11 probe is not oxidized by H_2_O_2_ itself, but by hydroxyl radicals formed in the Fenton reaction.

### Immunoprecipitation and LC-MS/MS

20 million HeLa cells expressing each target protein were washed once and harvested by centrifugation at 2000 rpm for 3min at 4°C. The cell pellet was suspended in RSB300 (10 mM Tris-HCl, pH 7.8, 300mM NaCl, and 2.5mM MgCl_2_) containing 1% Empigen BB, 1% DDM (n-dodecyl-ß-D-maltoside), 70mM sucrose,1x protease inhibitor cocktail (Roche) and 1x phosphatase inhibitor cocktail (Roche) and incubated on ice for 15min, the lysate was then homogenized with a 27-G needle (BD) for five times. 5% of the whole cell lysates was kept as input sample and the left was centrifuged at 15,000g for 30min at 4°C. The supernatant was incubated with anti-FLAG M2 affinity gel (Sigma) for 2h at 4°C. The gel was washed by RSB150 (10mM Tris-HCl, pH 7.8, 150mM NaCl, and 2.5mM MgCl_2_) containing 0.05% Triton X100 and 0.1% DDM three times and eluted by RSB300 containing 1% DDM and 250μg/mL 3xFLAG peptide. The eluates were added 1x LDS buffer and 100mM DTT as the final concentration, electrophoresis on 4-12% NuPAGE gel and stained by SimplyBlue SafeStain (Invitrogen). The entire stained gel lanes were excised were reduced with TCEP, alkylated with iodoacetamide, and digested with trypsin. Tryptic digests were analyzed using a standard 90min LC–MS/MS for IP samples and an extended 4h LC–MS/MS for Input samples on the Q Exactive Plus mass spectrometer (Thermo Fisher Scientific) with a nanoACQUITY Ultra Performance LC system (Waters).

### Proteomic data analysis

Raw data were analyzed by MaxQuant 1.6.3.3 using the UniProt Human Proteome database (08/21/2023) plus a contaminant database and protein-protein interactions were determined using label-free quantification. Variable modifications considered in the search were protein N-terminal acetylation, Asparagine deamidation, and methionine oxidation. Protein and peptide false discovery rate was set at 1%. The proteinGroup.txt file from MaxQuant was used as input matrix for Perseus 2.0.11 with the respective LFQ intensity as the main columns. Filter rows based on categorical column” to exclude proteins identified by site, matching to the reverse database or contaminants. Transform the intensity to a logarithmic scale and filter rows containing at least 1 valid value across all samples, missing values were imputed from normal distribution (width: 0.3, down shift: 1.8). Fold change and p-value were generated based on transformed LFQ intensities with the following settings: test, Student’s t-test; side, both; number of randomizations, 250; preserve grouping in randomizations, <NONE>; permutation-based FDR, 0.05; and s0, 0.1. Further bioinformatics analysis was conducted in Microsoft Excel and R statistical computing software (4.1.3). Significantly enriched co-IP proteins with ranked values (log2FC>1.0, p-value<0.05) in each sample were used to search for protein networks in STRINGdb (https://string-db.org/) v12.0 including each protein’s log2FC in functional enrichment analysis and with medium confidence (0.4) evidence from experimental interaction data, co-expression data, gene fusions, gene co-occurrence and gene neighborhood. In cases where the MS peptides do not distinguish between two or more proteins, only the first listed was used (e.g., RSP27A; UBA; UBC etc. only RPS27A was used). The resulting network were augmented by Inclusion of highest confidence STRING predicted functional partners (STRING’s built-in “More” selection) and sorted into functional clusters using built-in MCL clustering (natural clusters based on the stochastic flow) with inflation default parameters of 3. The pathway and process enrichment analyses were exported from STRING website.

## Acknowledgements

We thank the Penn Medicine Cell & Developmental Biology Microscopy Core for excellent help with confocal microscopy; the Wistar Institute Proteomics and Metabolomics Facility for expert assistance with the MS analysis; and the Pancreatic Islet Cell Biology Core for valuable advice on cellular metabolic measurements. This work was supported by a grant from the National Institutes of General Medical Sciences (R35 GM139646), and the Howard Hughes Medical Institute Investigator funding to G.D.

## Author contributions

J.X., J.D., Z.C., C.A., and G.D. conceived and designed the study. J.X., J.D., Z.C., C.A., J.X., M.J., and B.R.S. performed the experiments. J.D., J.X., C.D. and C.C.V performed the bioinformatics analysis. J.X., J.D., Z.C., and G.D. wrote the manuscript with input from all authors. G.D. is responsible for the project’s planning, supervision, experimental design, and funding acquisition.

## Competing interests

The authors declare that they have no competing interests.

## Data and materials availability

All data needed to evaluate the conclusions in the paper are present in the paper and/or the Supplementary Materials. RNA sequencing data have been made available on the NCBI Gene Expression Omnibus under accession GSE224868. The studies have also reanalyzed multiple datasets that are publicly available: GSE168831, GSE67528, GSE171343, GSE134737, GSE142499, GSE67196 and GSE163952.

## Supplementary Materials

**Fig. S1.**
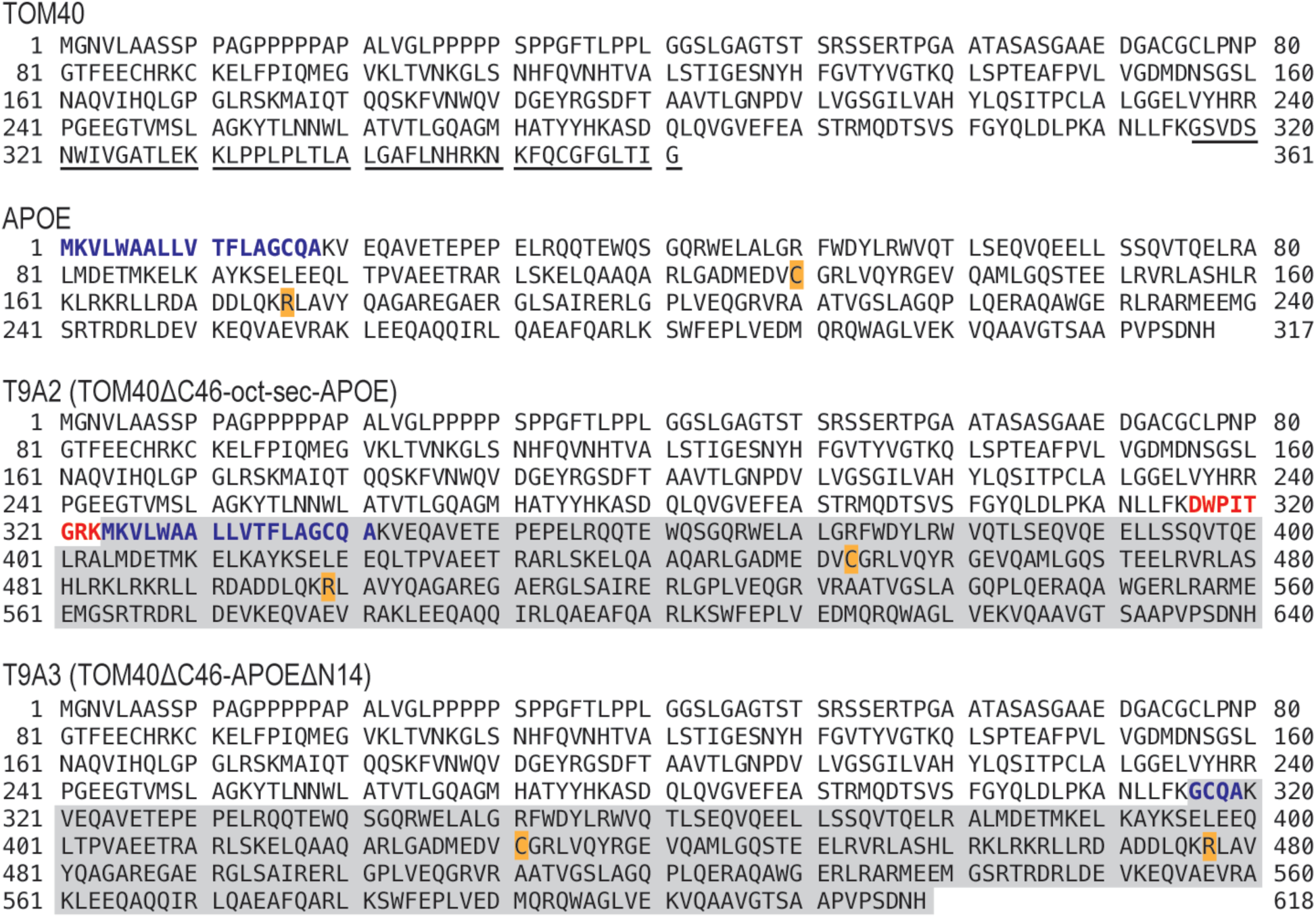
Amino acid sequences of human TOM40, APOE and the predicted chimeric protein, T9A2 (TOM40ΔC46-oct-sec-APOE) and T9A3 (TOM40ΔC46-APOEΔN14), formed by splicing of the TOMM40-APOE transcript. The TOM40 sequence shown here is the canonical isoform (uniprot O96008), MW = 38kDa, and the C-terminal 46aa encoded by TOMM40 exon10 (T10) are underlined. The APOE sequence shown here is the APOE3 isoform (uniprot P02649), MW = 36kDa, and the residues varied in other APOE isoforms are highlighted in orange. In detail, C130 and C176 in APOE2, R130 and R176 in APOE4. The same color scheme applies to the following sequences. The T9A2 sequence shown here is predicted, MW = 70kDa. The octapeptide (shown in bold red) predicted to be encoded in-frame from the splicing of TOMM40 exon9 to the portion of the APOE 5‘UTR in exon2 followed by APOE 18aa N-terminal secretory signal (shown in bold blue). T9A2 lacks TOM40 C-terminal 46aa encoded by TOMM40 exon10.The T9A3 sequence shown here is predicted, MW = 68kDa. Compared to T9A2, T9A3 lacks 22aa including the 8 aa octapeptide and 14 out of 18aa of the secretory signal. The remaining 4aa of the secretory signal are shown in bold blue. The APOE portion in T9A2 or T9A3 is highlighted in gray.

**Fig. S2.**
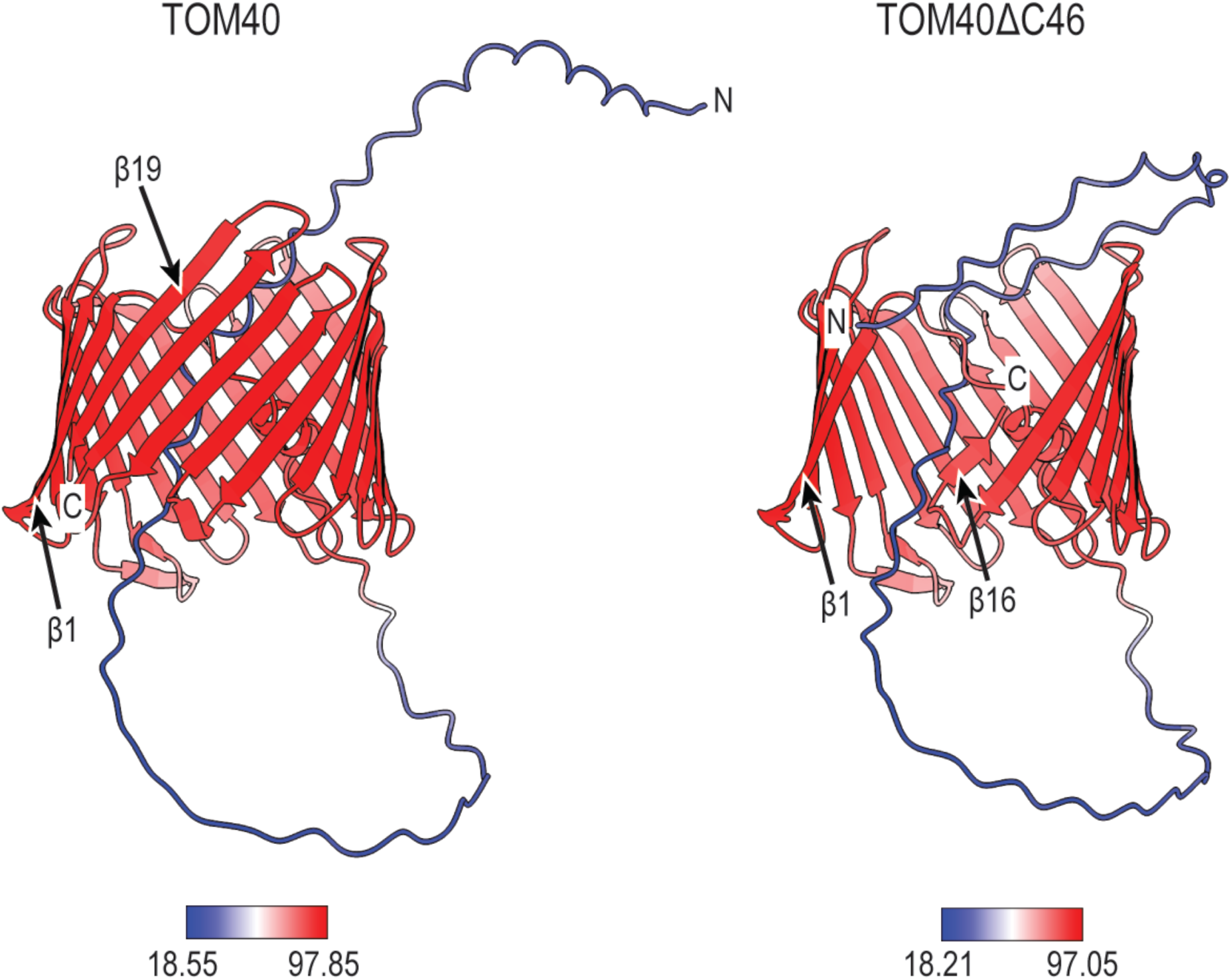
AlphaFold2 predicted TOM40 and TOM40ΔC46 structures. Overall view of AlphaFold2 predicted models of TOM40 showing very high confidence beta-barrel fold as in the experimentally determined structures. Note the very high similarity in the beta-barrel of TOM40ΔC46. The structures were rendered in ChimeraX and colorized by AlphaFold2 confidence metric (pLDDT), per-residue on the ribbon structure, as shown in deep red and blue, respectively. Minimum and maximum values for pLDDT are indicated.

**Fig. S3.**
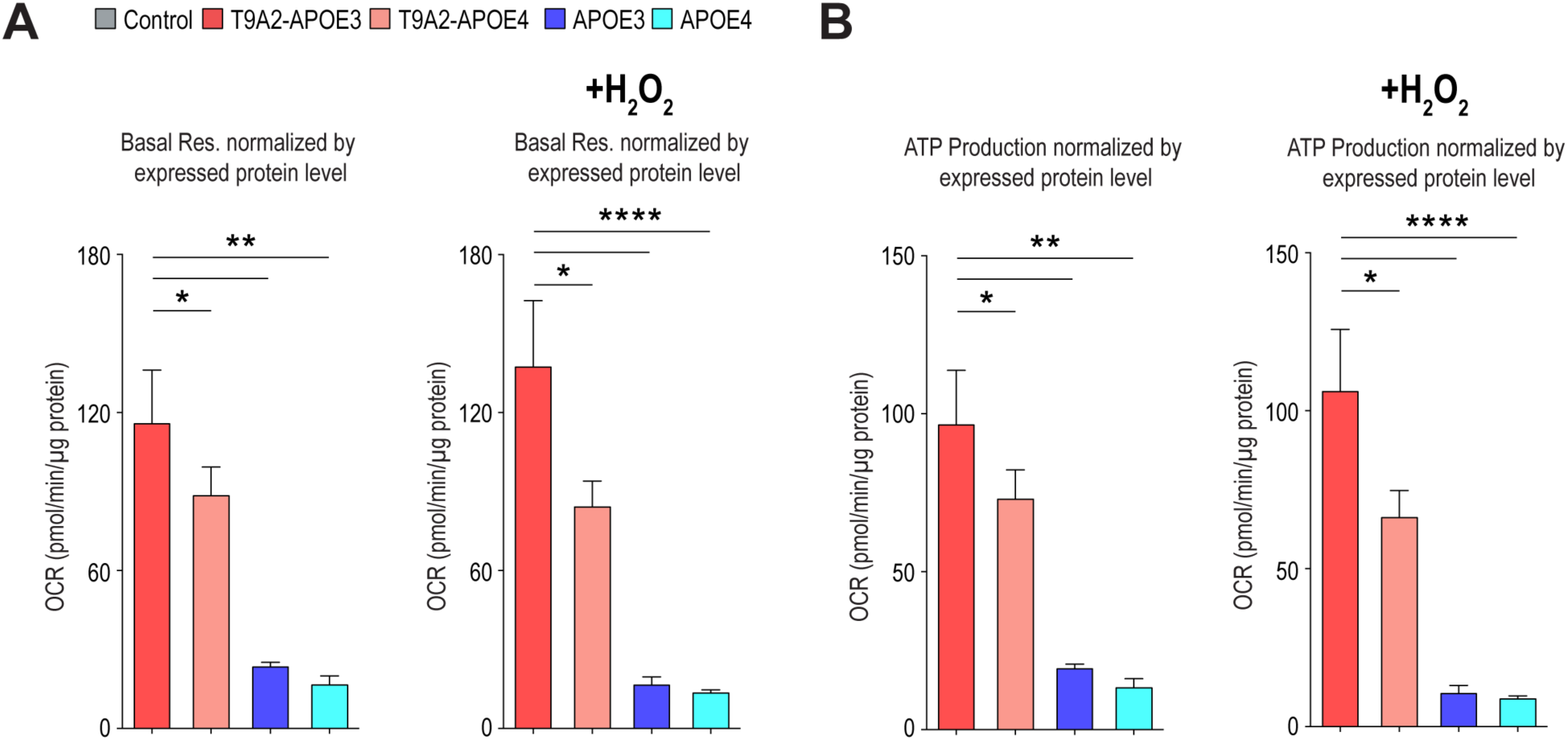
T9A2s strongly boost mitochondria respiration compared with APOEs. (**A**) Basal respiration normalized to the corresponding expressed protein levels determined from western blots. (**B**) ATP production normalized to the corresponding expressed protein levels determined from western blots. One-way ANOVA analyses were performed for the p-value. (*P < 0.05, **P < 0.01, ***P < 0.001 and ****P < 0.001).

**Fig. S4.**
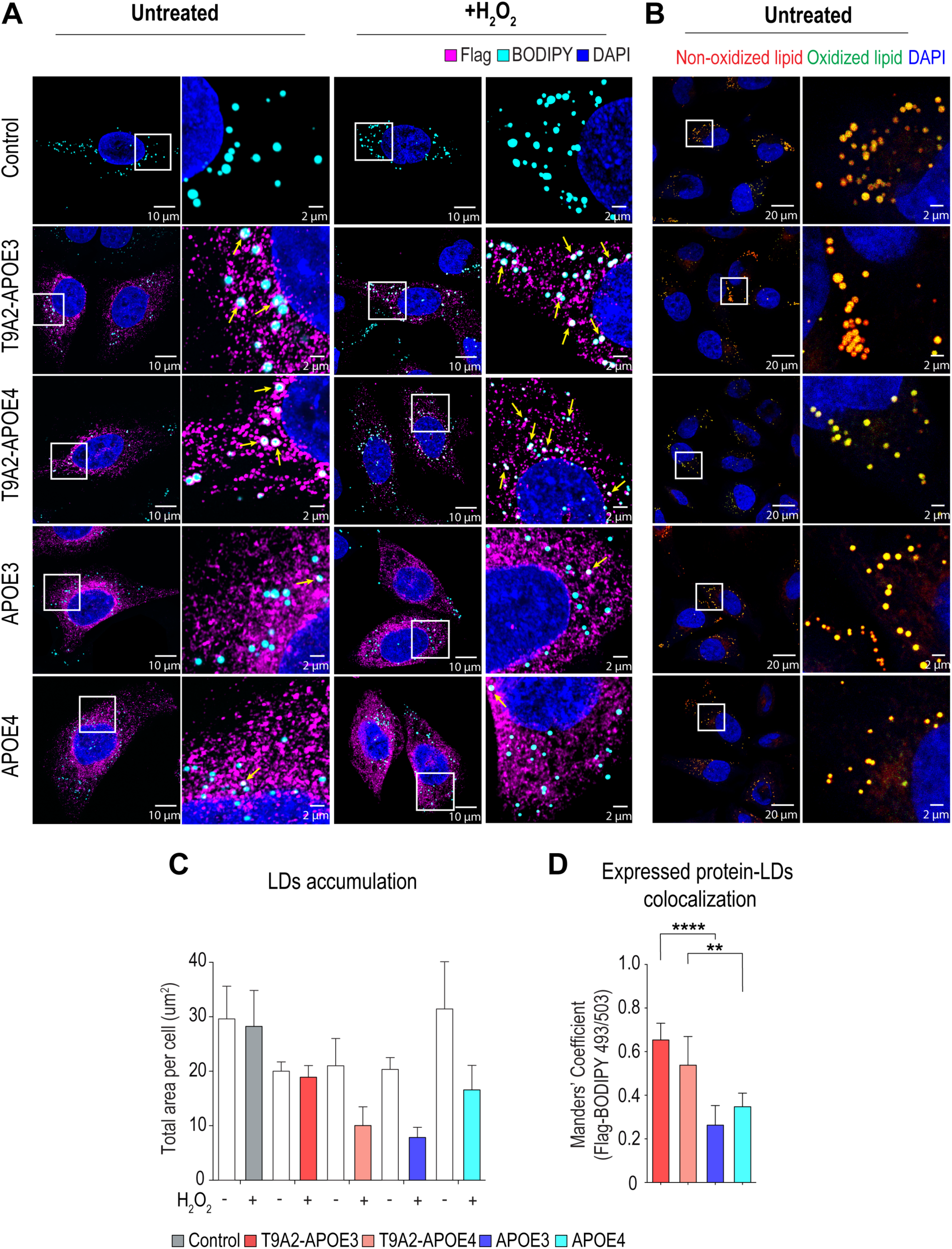
T9A2-APOE3 promotes lipid droplet formation under oxidative stress. (**A**) Representative images of HeLa CCL2 cells expressing Flag-tagged T9A2-APOE variants (3 and 4) and the corresponding APOE variants at 18h post-DOX induction, following with 100μM H2O2 treatment for 1h to induce oxidative stress. The cells are stained with BODIPY 493/503 (neutral lipid, cyan), anti-Flag (target protein, magenta), and DAPI (blue). Yellow arrows indicate co-localization of BODIPY stained LD and Flag-tagged protein in the same optical sections. Scale bar, 10μm and 2μm. (**B**) Representative images of BODIPY 581/591 C11 staining without H_2_O_2_ treatment. Non-oxidized lipid (red), and oxidized lipid (green) staining are shown indicated. Scale bar, 20μm and 2μm. (**C**) Quantification of the amount (total area) of lipid droplets per cell. n=15-18. (**D**) Histogram showing the colocalization (Mander’s coefficient) of the Flag and BODIPY 493/503 signals in HeLa cells. n=15-18

**Fig. S5.**
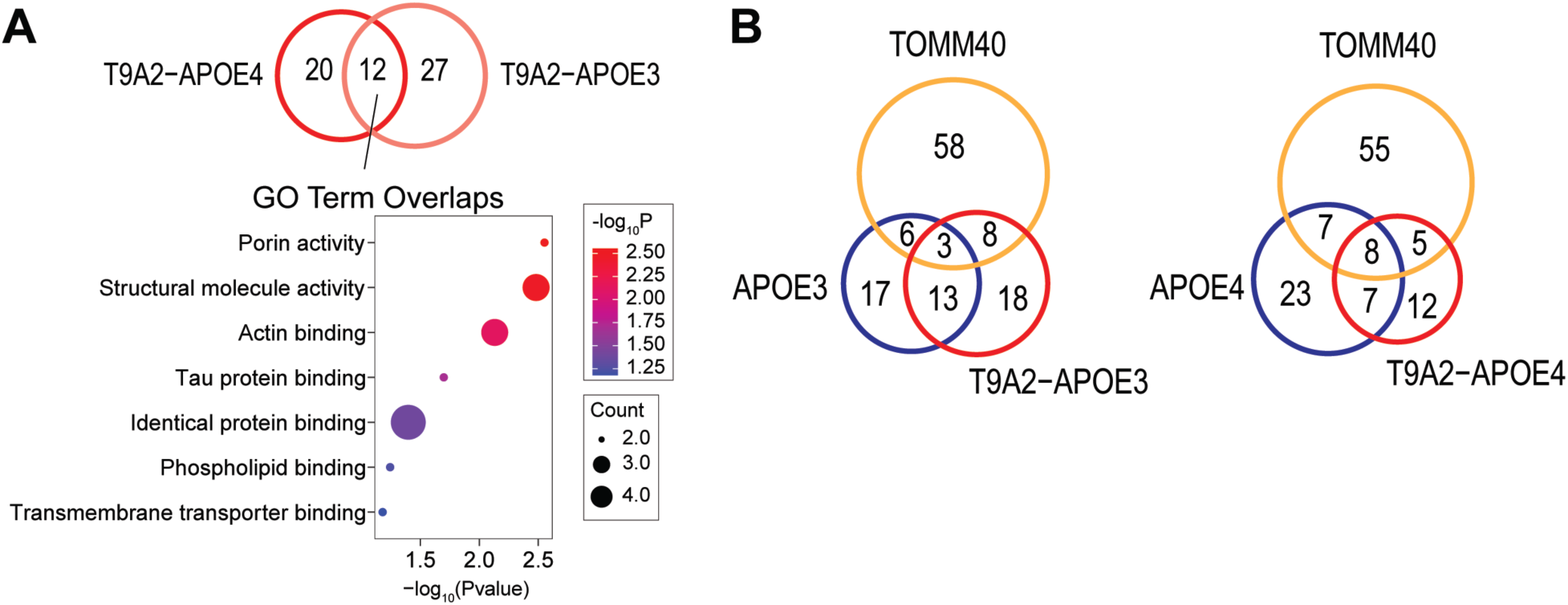
T9A2’s interactome profiling. (**A**) Top: Venn diagram of significantly enriched proteins (|log2FC| >1, -log10(p-value) >1.3) in T9A2-APOE3 and T9A2-APOE4. Bottom: GO term analysis of common interactors from T9A2-APOE3 and T9A2-APOE4. (**B**) Venn diagram depicting the overlap of significantly enriched proteins in T9A2-APOE3 and T9A2-APOE4 compared with the corresponding APOE variants and TOMM40.

**Fig. S6.**
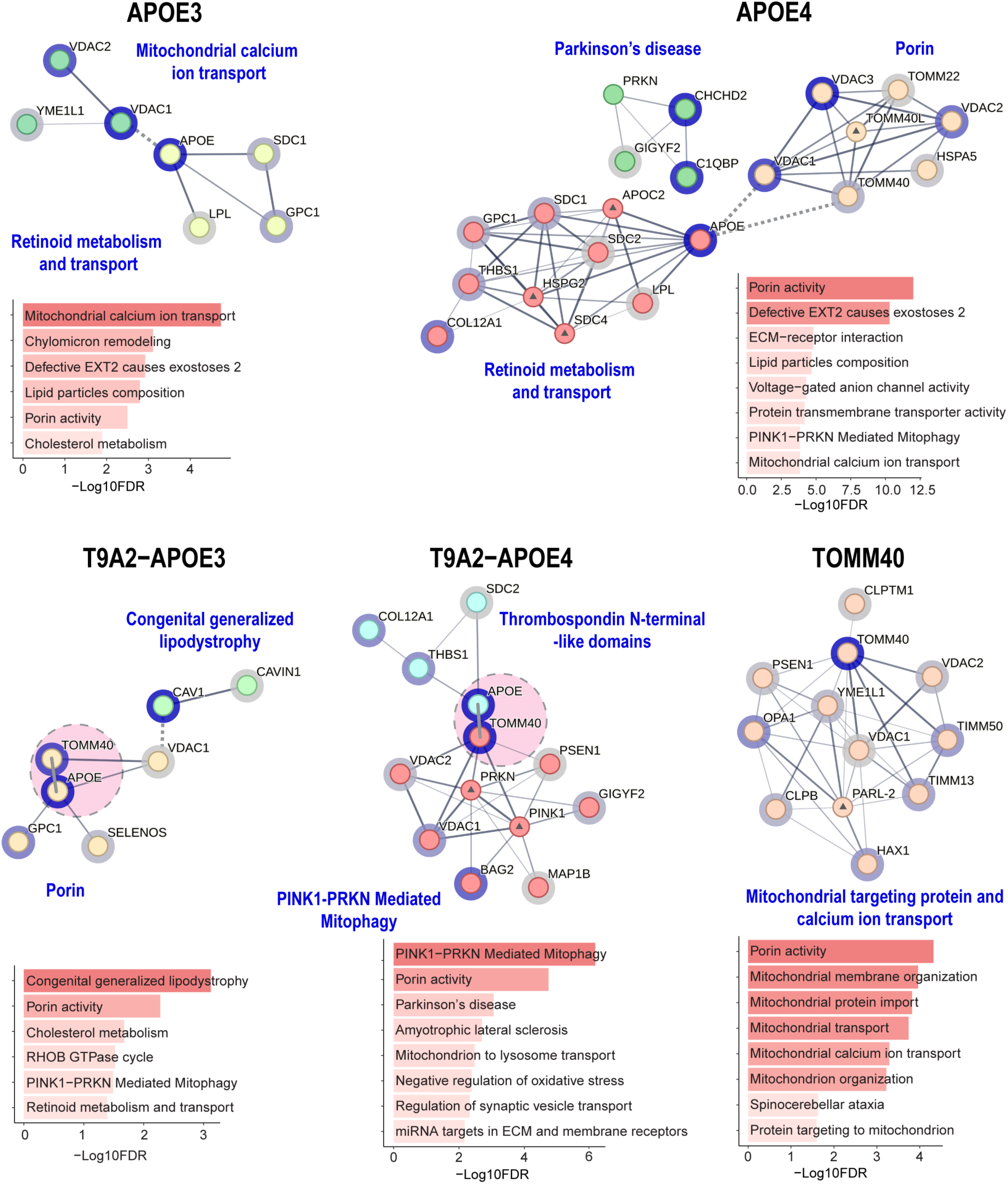
T9A2’s, APOE’s and TOMM40’s interaction partners regulate different cellular pathways. Protein–protein interaction networks of the proteins detected in IP-MS of the indicated Flag-tagged T9A2s, APOE, and TOMM40 from HeLa cells, analyzed for each along with their fold enrichment vs control in duplicates (log2FC>1.0; p-value<0.05) and visualized in STRING. The main functional clusters (MCL clustering, (FDR=<0.05) that contain APOE and/or TOMM40 are marked on each network. Each protein is encircled in blue of intensity corresponding to their abundance (log2FC enrichment rank) in that IP. The nodes with a central triangle represent predicted functional partners from the STRING database. APOE and TOMM40 in T9A2-APOE3 and T9A2-APOE4 are shown as touching with their centers connected by a short thick line to indicate they are in fact one protein and highlighted in an ellipsoid. Analyses of enriched proteins in selected clusters from each IP-MS (log2FC1.0; p-value<0.05) by Gene Ontology (GO), KEGG, Reactome, and WikiPathways, are shown under each network. FDR, false discovery rate.

**Table S1.**
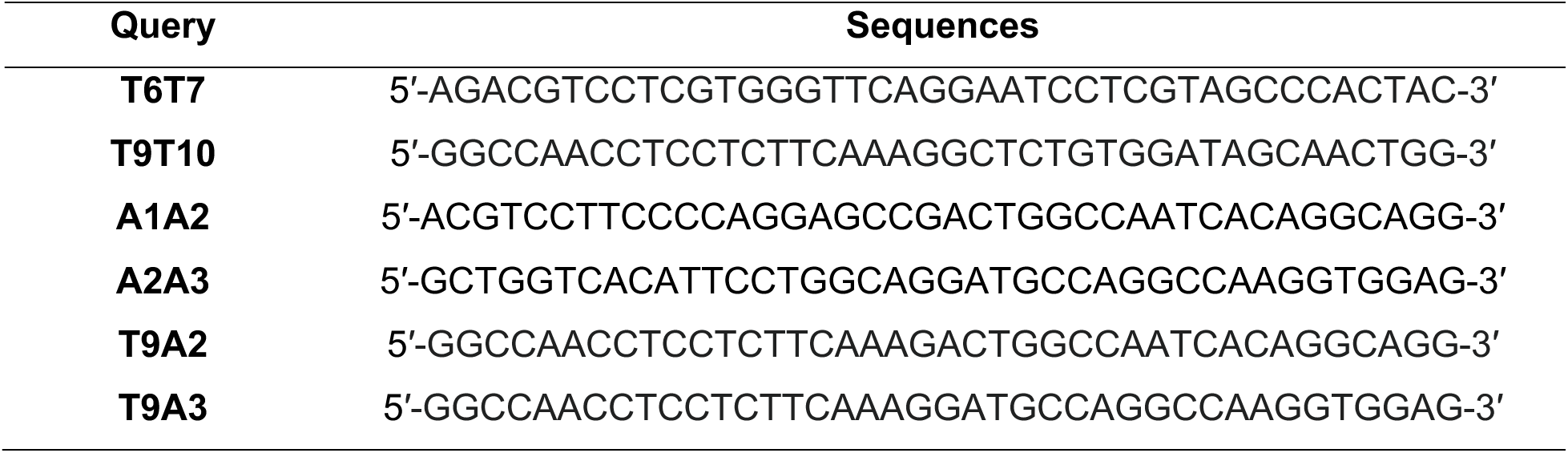
Query sequences used in BigBlast.

## Notes

### Competing Interest Statement

The authors have declared no competing interest.

